# Coancestry superposed on admixed populations yields measures of relatedness at individual-level resolution

**DOI:** 10.1101/2024.12.29.630632

**Authors:** Danfeng Chen, John D. Storey

## Abstract

The admixture model is widely applied to estimate and interpret population structure among individuals. Here we consider a “standard admixture” model that assumes the admixed populations are unrelated and also a generalized model, where the admixed populations themselves are related via coancestry (or covariance) of allele frequencies. The generalized model yields a potentially more realistic and substantially more flexible model that we call “super admixture”. This super admixture model provides a one-to-one mapping in terms of probability moments with the population-level kinship model, the latter of which is a general model of genome-wide relatedness and structure based on identity-by-descent. We introduce a method to estimate the super admixture model that is based on method of moments, does not rely on likelihoods, is computationally efficient, and scales to massive sample sizes. We apply the method to several human data sets and show that the admixed populations are indeed substantially related, implying the proposed method captures a new and important component of evolutionary history and structure in the admixture model. We show that the fitted super admixture model estimates relatedness between all pairs of individuals at a resolution similar to the kinship model. The super admixture model therefore provides a tractable, forward generating probabilistic model of complex structure and relatedness that should be useful in a variety of scenarios.

## 1 Introduction

Populations are structured when genotype frequencies do not follow Hardy-Weinberg proportions. This may be due to several factors, including finite population sizes, migration, and genetic drift [1, 2]. Our goal here is to develop a framework and estimation method of a forward generating probability process that captures the observed genetic structure and relatedness among a set of individuals in a population-based study.

The framework is based on covarying allele frequencies among populations [3] and individuals [4], which we will refer to as *coancestry* [3–5]. The data underlying the proposed method are single nucleotide polymorphism (SNP) genotypes measured throughout the genome on a set of individuals. The aim is to formulate and estimate a model of the underlying process that leads to individual-specific allele frequencies (IAFs), which are parameters consisting of possibly distinct allele frequencies for every individual-SNP pair. IAFs have been formulated in previous work [6, 7] and they are the estimation target in several established admixture methods [8–10], a genome-wide association test for structured populations [11], and a test of structural Hardy-Weinberg equilibrium [12].

A joint probability distribution of the IAFs under a neutral model has been developed that yields covariances for all pairs of IAFs, parameterized by ancestral allele frequencies and coancestry parameters [4, 5]. This model produces a one-to-one mapping with the kinship parameters from the *identity-by-descent* model [13, 14], excluding close familial genetic relationships. This coancestry model therefore captures pairwise individual-level structure and relatedness equivalent to the kinship model. However, similarly to the kinship model, the coancestry parameterization is in terms of expected values, variances, and covariances of the IAFs and genotypes. It does not explicitly define a forward-generating probability model of IAFs.

Admixture models have been explored as a possible way to define such a forward-generating probability model [4, 5]. The products of an admixture model are individual-specific admixture proportions and population-specific allele frequencies. The IAFs are modeled as a weighted average of these *antecedent population allele frequencies* by the *individual-specific admixture proportions*. Several methods treat the admixture proportions and antecedent population allele frequencies as unknown parameters without explicitly making any assumptions about their random distributions [8–10]. Other methods place a prior probability distribution on them for Bayesian modeling fitting purposes [15–17]; however, these Bayesian methods do not include these prior distributions as an inference target.

In considering a model of random antecedent population allele frequencies, one could assume that the allele frequencies are independently generated among all antecedent populations based on a common set of parameters (e.g., independent draws from the Balding-Nichols distribution [18]). We will call this assumption the “standard admixture” model. However, this standard admixture model may be overly restrictive; rather, one could implement a coancestry model of the antecedent allele frequencies according to pairwise covariances [4, 5]. We will call this model the “super admixture” model, as coancestry (or covariance) is superposed on the admixed antecedent populations. Fig. 1 displays a schematic of these models.

**Figure 1:**
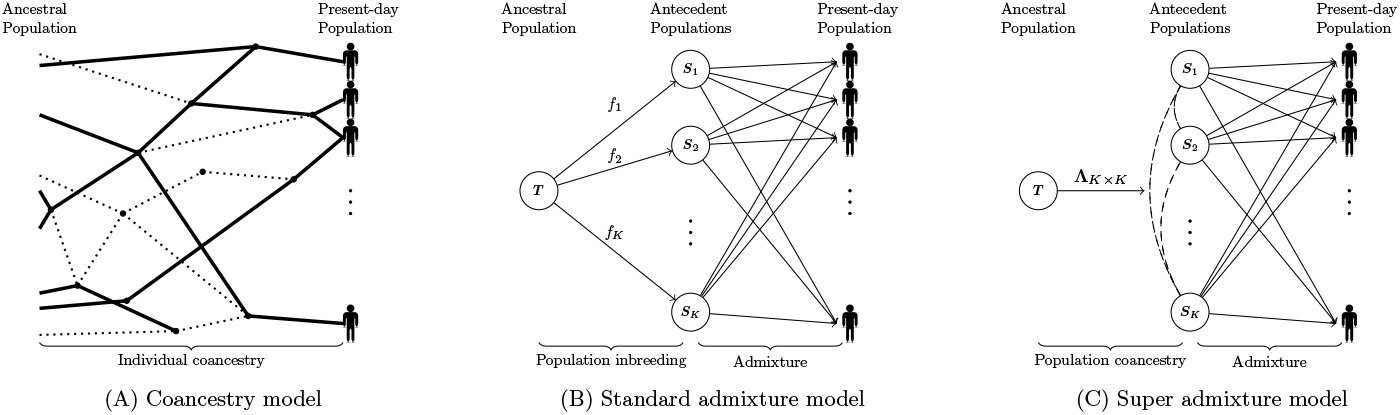
Graphical representations of the coancestry model, the standard admixture model, and the super admixture model. (A) In the coancestry model, individuals in the present-day population are connected by a complex genealogy. (B) In the standard admixture model, the arrows connecting *T* with *S*_1_, … , *S*_*K*_ reflect that the antecedent populations evolved independently from *T* . Arrows connecting *S*_1_, … , *S*_*K*_ with individuals in the present-day population reflect that these individuals were admixed from independent antecedent populations. (C) In the super admixture model, dashed lines connecting all pairs of antecedent populations reflect that antecedent populations have coancestry parameterized by **Λ.** Arrows connecting *S*_1_, … , *S*_*K*_ with individuals in the present-day population reflect that these individuals were admixed from covarying antecedent populations.

Here, we develop a method that estimates the parameters in the super admixture model, which includes the standard admixture model as a special case. The method is based on method of moments estimation and geometric considerations, so it does not make assumptions about the probability distributions of the parameters and it does not involve costly likelihood maximization computations. Likelihood maximization is the most common approach used in fitting the admixture models [8, 9, 15–17], but we build from a recently proposed distribution-free moment-based method, called ALStructure, that only uses linear projections and geometric constraints on parameters to estimate the model [10]. ALStructure performs favorably to likelihood based methods (even in achieving a high likelihood) and can be tractably scaled to massive data sets. Our proposed super admixture method complements this framework and has similar advantages.

We establish super admixture through computational studies and analyses of data sets, including the human genome diversity panel (HGDP) [19], the 1000 genomes project (TGP) [20], the Human Origins study (HO) [21], and a study on individuals with Inadian ancestry (IND) [22]. We show on all of these data sets that the super admixture method is capable of capturing the same relatedness and structure as a model-free individual-level coancestry estimator [4], whereas the standard admixture model does not. We demonstrate that the framework can generate bootstrap genotypes that retain the structure seen in the human studies. For example, Fig. 2 shows these results on the HO study. We show that the coancestry among antecedent populations estimated by super admixture yields new insights and visualizations of structure previously unavailable, for example, Fig. 3 on the HO study. We develop and perform a statistical test to demonstrate on the studies that coancestry among the admixed antecedent populations is statistically different from zero to an high degree of significance.

**Figure 2:**
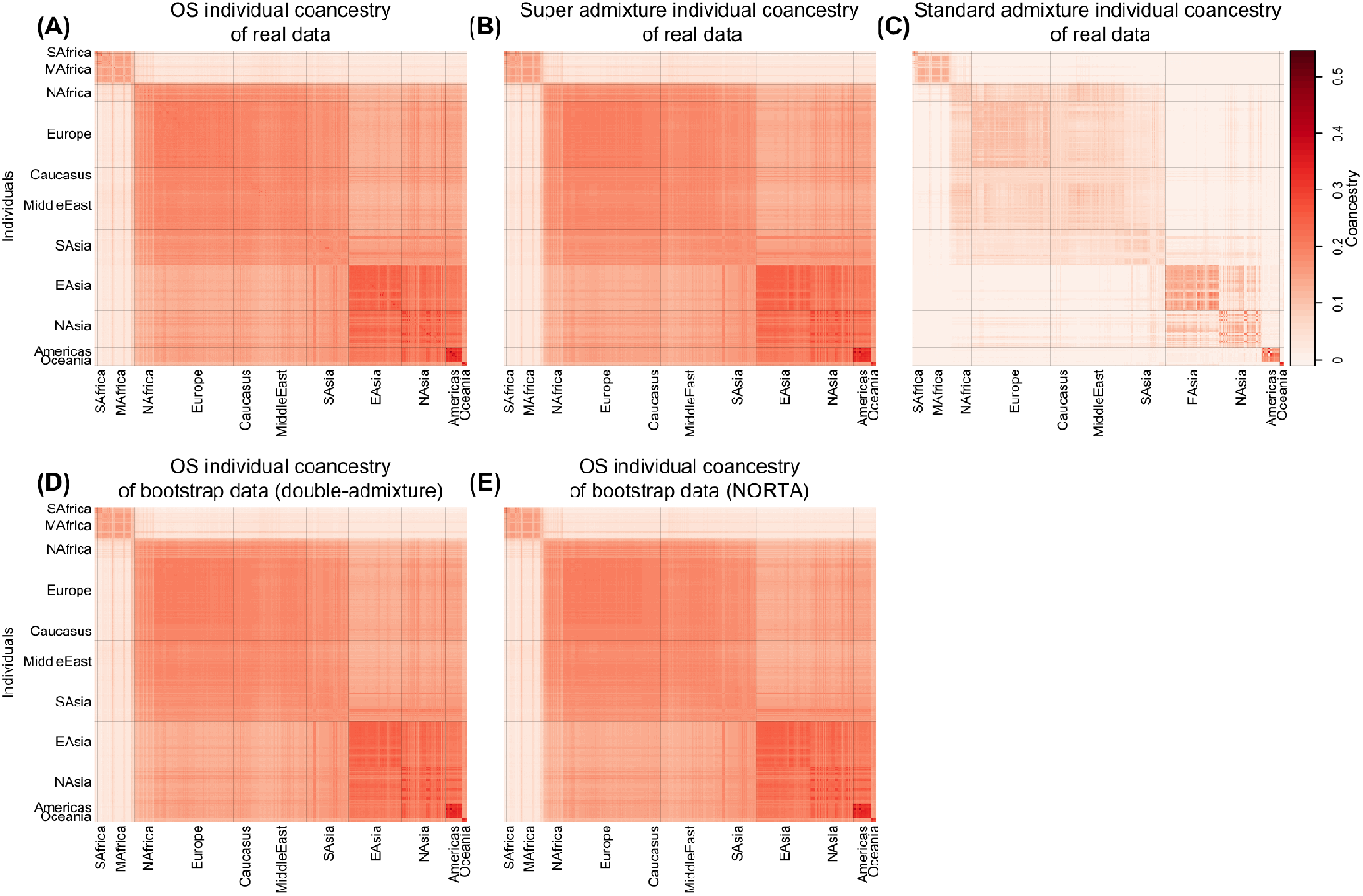
Heatmaps of individual-level coancestry estimates in the HO data set.

**Figure 3:**
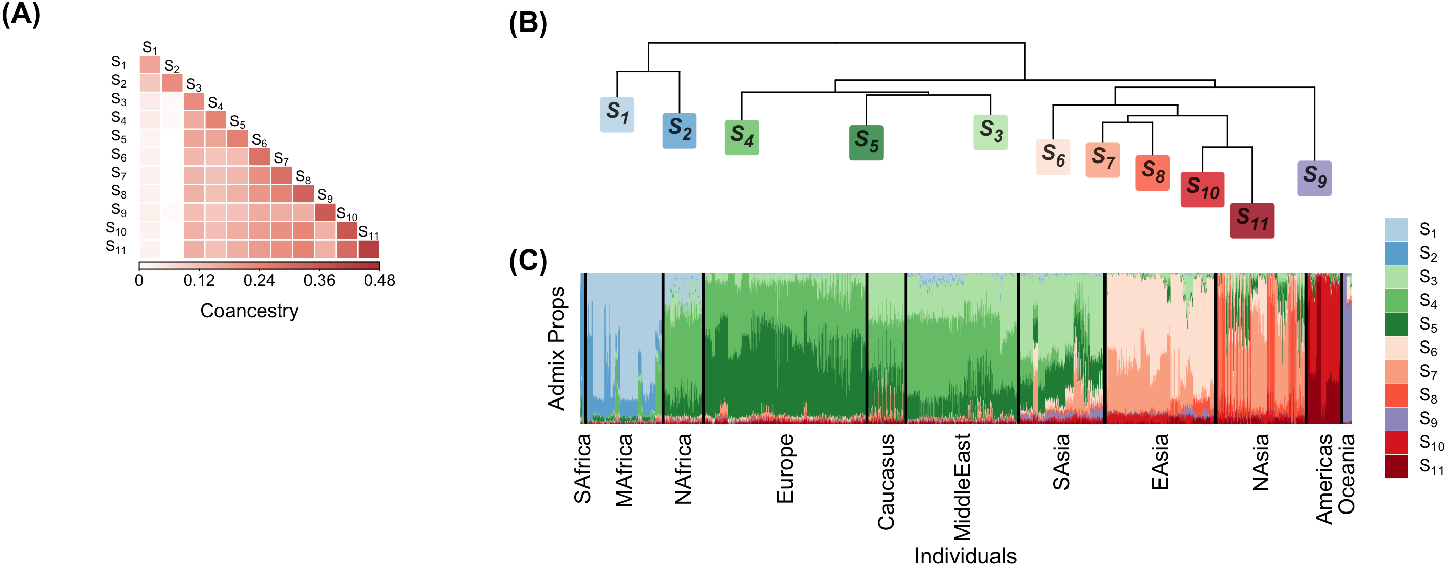
(A) Heatmap of antecedent population coancestry estimates in the HO data set. (B) Dendrogram representation of the antecedent population coancestry estimates. (C) Stacked bar plot of admixture proportions.

Our proposed framework makes several contributions: (i) a distribution-free framework that can account for arbitrarily complex relationships among the admixed antecedent populations in the admixture model; (ii) admixture-based estimation of individual-level pairwise coancestry at a resolution equivalent to general, model-free coancestry and kinship; (iii) a partitioning of the super admixture model into evolutionary, genealogical, and statistical sampling components; and (iv) a tractable algorithm to form bootstrap samples of genotypes from the estimated evolutionary process.

## 2 Super admixture framework

Here, we first introduce the data and models, and then we detail the proposed framework. We describe how the framework is used to estimate the super admixture model, generate parameters and data from the model, and perform a hypothesis test of the standard versus super admixture models.

### 2.1 Coancestry

We assume that *m* SNPs are measured on *n* individuals. The genotype measurements are denoted by *x*_*ij*_ for *i* = 1, … , *m* and *j* = 1, … , *n*. For each SNP, one of the alleles is counted as a 0 and the other as a 1, implying that the SNP genotypes are *x*_*ij*_ ∈ {0, 1, 2} where *x*_*ij*_ = 0 is homozygous for the 0 allele, *x*_*ij*_ = 1 is a heterozygote, and *x*_*ij*_ = 2 is homozygous for the 1 allele. We assume that 𝔼[*x*_*ij*_|*π*_*ij*_] = 2*π*_*ij*_ for IAF *π*_*ij*_. This IAF parameterization allows each individual-SNP pair to possibly have a distinct allele frequency. The classical scenario where there is one allele frequency per SNP is a special case where *π*_*i*1_ = *π*_*i*2_ = · · · = *π*_*in*_. The conditional expected value 𝔼[*x*_*ij*_|*π*_*ij*_] = 2*π*_*ij*_ also allows for the IAFs *π*_*ij*_ to be random parameters, which we assume here.

We utilize an existing coancestry model where the IAFs are random parameters with respect to some ancestral population *T* that is common to all *n* individuals [4, 5]. This is a neutral model where

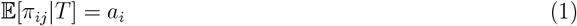

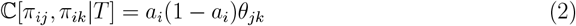

for *i* = 1, … , *m* and *j, k* = 1, … , *n*.The parameter *a*_*i*_ is the ancestral allele frequency in *T* for SNP *i* and 0 ≤ *θ*_*jk*_ ≤ 1 is the coancestry for individuals *j* and *k* with respect to *T* . (Note that the *a*_*i*_ and *θ*_*jk*_ parameters depend on *T* and could be different if conditioning on a different common ancestral population.) The coancestry model we utilize also makes the assumption used in previous work [4, 5, 7–12] that

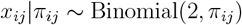

where the *x*_*ij*_ are jointly independent. Under this model, it follows that

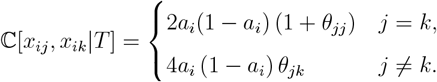

A one-to-one mapping exists with the identity-by-descent kinship model (often used in GWAS methods), denoted by *ϕ*_*jk*_, by matching variances and covariances [4, 5]. The parameters map so that

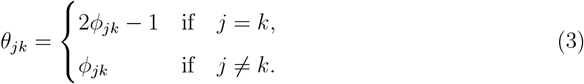

When min_*jk*_ *θ*_*jk*_ = 0, then *T* is the most recent common ancestral population [4]. The full set of parameters is denoted by the *n* × *n* symmetric matrix **Θ** with (*j, k*) entry *θ*_*jk*_.

### 2.2 Admixture models

#### General admixture

We first describe a general formulation of the admixture model, of which standard and super admixture are special cases. There are *K* populations *S*_1_, *S*_2_, … , *S*_*K*_ descended from *T* that precede the present day population, which we refer to as “antecedent populations”. While *T* has allele frequencies *a*_1_, *a*_2_, … , *a*_*m*_, antecedent population *S*_*u*_ has allele frequencies *p*_1*u*_, *p*_2*u*_, … , *p*_*mu*_ for *u* = 1, 2, … , *K*. The allele frequencies {*p*_*iu*_} are random parameters from a distribution parameterized by {*a*_*i*_} plus other possible parameters that characterize the evolutionary process from *T* to *S*_*u*_.

For each individual *j*, there is a genealogical process from population *T* to the present day population. This is captured by a random *K*-vector *q*_1*j*_, *q*_2*j*_, … , *q*_*Kj*_ of admixture proportions, where 0 ≤ *q*_*uj*_ ≤ 1 and 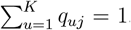. The parameter *q*_*uj*_ is the proportion of the individual *j* randomly descended from *S*_*u*_. Therefore, the IAFs are such that

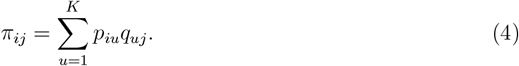

We collect the antecedent population allele frequencies into the *m* × *K* matrix ***P*** and the admixture proportions into the *K* × *n* matrix ***Q***, it follows that

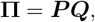

where **Π** is an *m* × *n* matrix with (*i, j*) entry *π*_*ij*_.

#### Standard admixture

We define the standard admixture model to be the case where the antecedent allele frequencies are independently distributed. Specifically, in this model *p*_*iu*_ is a random parameter with mean *a*_*i*_ and variance *a*_*i*_(1 − *a*_*i*_)*f*_*u*_. The standard admixture model is defined as follows for *i* = 1, 2, … , *m* and *u* = 1, 2, … , *K*.

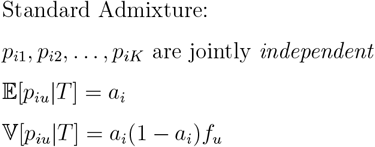

Under this parameterization, *a*_*i*_ is the ancestral allele frequency in *T* and *f*_*u*_ is the inbreeding coefficient or *F*_ST_ of antecedent population *S*_*u*_ with respect to *T* . Since the {*p*_*iu*_} are jointly independent, there is no coancestry among antecedent populations and there is no dependence among loci.

One well-known distribution that could be utilized here is the Balding-Nichols (BN) distribution [18] with parameters *a*_*i*_ and *f*_*u*_:

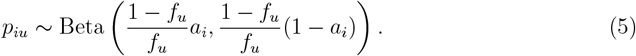

We will write this re-parameterized Beta distribution as BN(*a*_*i*_, *f*_*u*_). This achieves the expected value and variance of the standard admixture definition. The Balding-Nichols distribution is often used to generate allele frequencies for a set of populations to achieve desired expected allele frequencies and *F*_ST_ values. This distribution has been discussed as useful for generating antecedent allele frequencies in the standard admixture model [4–7, 23].

#### Super admixture

The super admixture model extends the standard admixture model in that it includes a covariance among antecedent population allele frequencies, which we refer to as population-level coancestry. While we denoted individual-level coancestry by *θ*_*jk*_, we will denote population-level coancestry by *λ*_*uv*_ for *u, v* = 1, 2, … , *K* where 0 ≤ *λ*_*uv*_ ≤ 1. We collect these values into the *K* × *K* symmetric coancestry matrix **Λ**. The super admixture model is defined as follows for *i* = 1, 2, … , *m* and *u, v* = 1, 2, … , *K*.

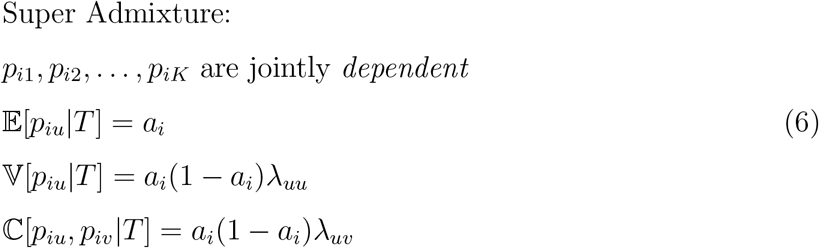

In this model we assume that allele frequencies between loci are independent, so the random *K*-vectors (*p*_*h*1_, *p*_*h*2_, … , *p*_*hK*_) and (*p*_*i*1_, *p*_*i*2_, … , *p*_*iK*_) are independent for *h* ≠ *i*. Thus, a potential generalization of the super admixture model is to include dependence among loci. Otherwise, the super admixture model is general in that it allows for the full range of coancestry values among antecedent populations.

#### Forward generating probability process

We now describe the super admixture model as a forward generating probability process. Suppose that the admixture proportions ***Q*** are drawn from some probability distribution 𝒬. Then, for *i* = 1, 2, … , *m* and *j* = 1, 2, … , *n*:

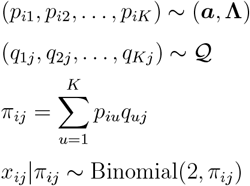

The joint probability of all random quantities can be factored as follows:

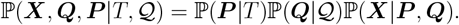

One interpretation of this is that ℙ(***P*** |*T* ) represents evolutionary sampling, ℙ(***Q***|𝒬) represents genealogical sampling, and ℙ(***X***|***P*** , ***Q***) represents statistical sampling.

#### Individual-level coancestry in the admixture models

Recall that in the covariance model, the covariance of two IAFs for a given SNP is ℂ[*π*_*ij*_, *π*_*ik*_|*T* ] = *a*_*i*_(1 − *a*_*i*_)*θ*_*jk*_, shown in Eq. (2). Conditioning on the admixture proportions ***Q***, which are ancillary to allele frequencies, this covariance under the super admixture model is, for *j, k* = 1, 2, … , *n*,

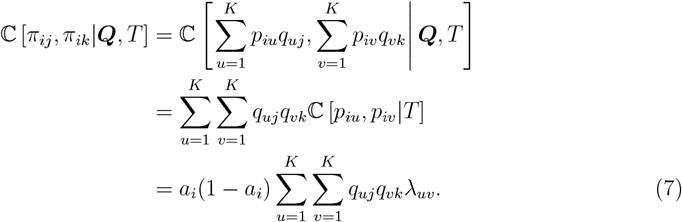

By setting the covariance from Eq. (2) equal to Eq. (7), it follows that under the super admixture model the individual-level coancestry is the following.

Super Admixture Individual-level Coancestry:

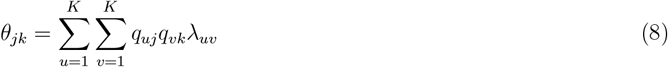

In the standard admixture model, 𝕍[*p*_*iu*_|*T* ] = *a*_*i*_(1−*a*_*i*_)*f*_*u*_, whereas in the super admixture model 𝕍[*p*_*iu*_|*T* ] = *a*_*i*_(1 − *a*_*i*_)*λ*_*uu*_. If we set *f*_*u*_ = *λ*_*uu*_, the difference between the standard and super admixture models is therefore that in the standard model, *λ*_*uv*_ = 0 for *u* ≠ *v*. To work with a single notation, we will therefore write *λ*_*uu*_ in place of *f*_*u*_ for the standard admixture model. The coancestry in this model is as follows.

Standard Admixture Individual-level Coancestry:

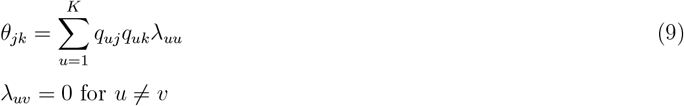

Considering all pairs of individuals simultaneously, the individual-level coancestry matrix **Θ** can be written in terms of the antecedent population-level coancestry **Λ** and the admixture proportions ***Q*** as

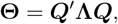

which is an important relationship we utilize to estimate **Λ**.

### 2.3 Estimating coancestry among antecedent populations

Here, we propose a method to estimate the antecedent population-level coancestry **Λ** under the super admixture model, with the standard admixture model estimate as a special case. The rationale is to leverage the relationship, **Θ** = ***Q***′**Λ*Q***. Given values for **Θ** and ***Q***, we identify values of **Λ** that make ***Q***′**Λ*Q*** close to **Θ**, while obeying the geometic constraints of **Λ** (i.e., 0 ≤ *λ*_*uv*_ ≤ 1 and *λ*_*uv*_ = *λ*_*vu*_).

Given values for **Θ** and ***Q***, we formulate the problem of the estimating the antecedent population-level coancestry **Λ** under the super admixture model as follows.

#### Problem 1

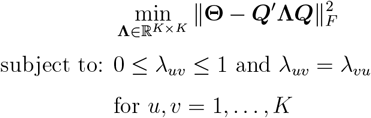

where ∥ · ∥_*F*_ represents the Frobenius norm defined in Appendix A.1. We utilize the proximal forward-backward (PFB) method [24] to solve this optimization problem, resulting in Algorithm 1 for solving Problem 1. Every sequence of (**Λ**_*t*_)_*t*∈ℕ_ generated from this algorithm is guaranteed to converge to a solution of the corresponding problem. The PFB method and how to employ it to our setting are detailed in Appendix B.2. The performance of Algorithm 1 is demonstrated in Appendix C.

##### Algorithm 1

Estimating **Λ** for the super admixture model given **Θ** and ***Q***

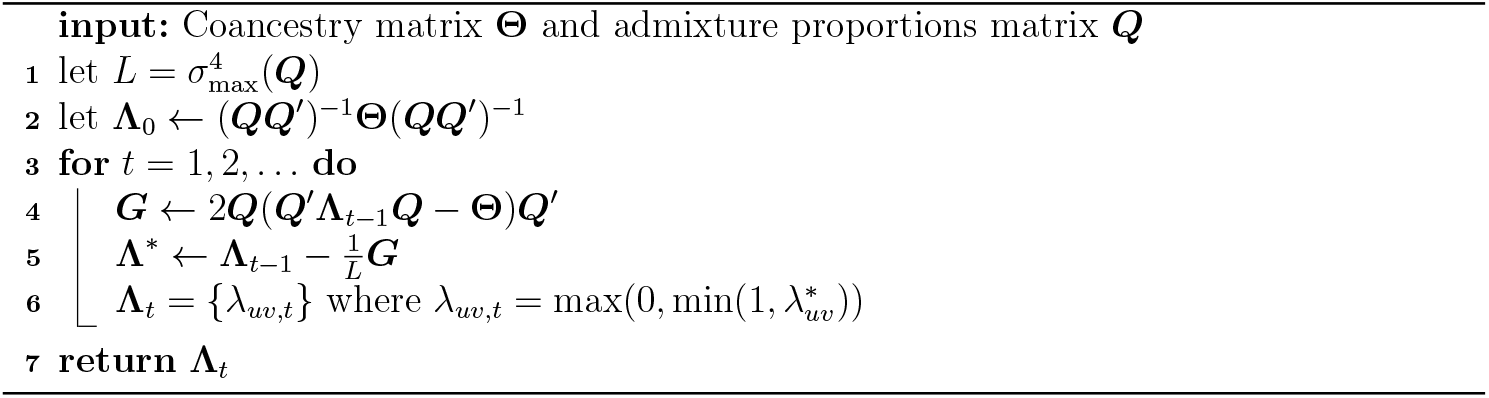

*σ*_max_(·) denotes the maximum singular value (Appendix A.1).

To estimate all components of the super admixture model, one needs estimates of the *n*× *n* individual-level coancestry matrix **Θ**, the *K*× *K* antecedent population-level coancestry matrix **Λ**, the *m* × *K* matrix of antecedent population allele frequencies ***P*** , and the *K* × *n* matrix of admixture proportions ***Q***. There exists a wide range of methods for estimating ***P*** and ***Q*** [9, 10, 15–17, 25]. Here, we utilize the ALStructure method [10], which implements method of moments and geometric constraints to estimate ***Q*** similarly to our approach here. In that method, a linear basis of ***Q*** is determined from ***X*** that has theoretical guarantees to span the true basis as the number of SNPs *m* grows large. A projection-based estimate 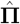 of the IAFs is also formed. The quantity 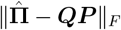 is then algorithmically minimized through geometrically constrained alternating least squares to yield estimates 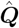 and 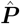.

We utilize the structural Hardy-Weinberg (sHWE) test [12] for determining the number of antecedent populations *K*, as outlined in that work. The approach is to consider a range of *K* values to test the assumption that *x*_*ij*_|*π*_*ij*_ ∼ Binomial(2, *π*_*ij*_) based on the estimates 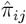 and a goodness-of-fit statistic with a parametric bootstrap null distribution; *K* is then parsimoniously chosen to satisfy this modeling assumption from a genome-wide perspective. A method of moments estimator of **Θ** was derived in ref. [4], where it was shown to have favorable properties and is consistent for the true values under certain assumptions. We denote this Ochoa-Storey (OS) estimate by 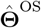 and review its details in Appendix B.1. If one has alternative ways to estimate **Θ** and ***Q***, and to determine *K*, then those can be used within our framework as well.

##### Algorithm 2

Estimating **Λ** for the super admixture model given ***X***

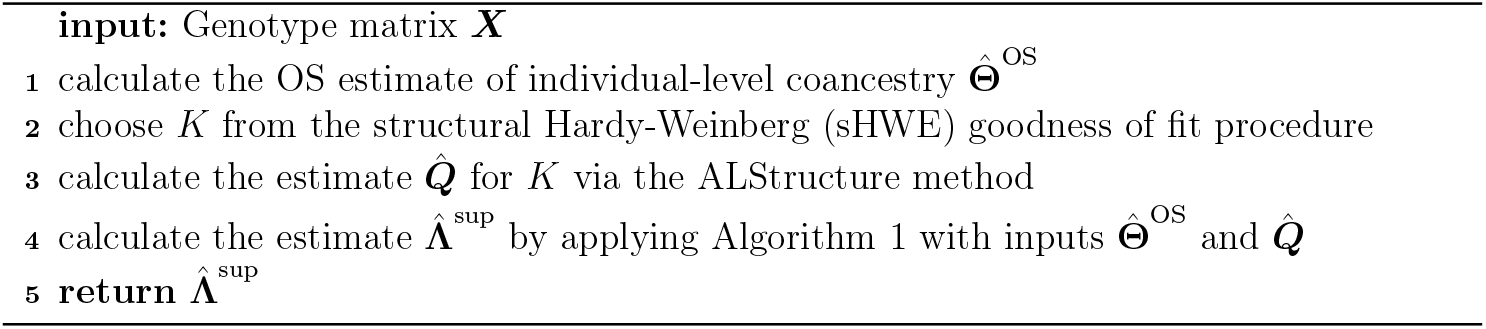

Note that one can further calculate a corresponding estimate for individual-level coancestry by

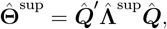

which can be compared to 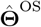 in order to aid in model fit assessment.

We can estimate **Λ** under the standard admixture model by modifying the constraints in Problem 1. This leads to Problem B.1 and Algorithm B.2 described in Appendix B.2. Algorithm 2 can then be used to form the estimate 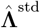 under the standard admixture model with Algorithm 1 in Line 4 replaced by Algorithm B.2. The corresponding estimate for individual-level coancestry can be calculated as 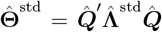. The performance of Algorithm B.2 is also demonstrated in Appendix C.

### 2.4 Simulating antecedent population allele frequencies

We now introduce a method to generate antecedent population allele frequencies with given coancestry **Λ**. We noted above in Eq. (5) that for the standard admixture model, one way to generate allele frequencies *p*_*i*1_, *p*_*i*2_, … , *p*_*iK*_ is via independent realizations from the Balding-Nichols (BN) distribution: *p*_*iu*_ ∼ BN(*a*_*i*_, *λ*_*uu*_) for *u* = 1, 2, … , *K*. As there is no default approach to extending this to the super admixture case, we propose a method here called “double-admixture”. The main idea of the method is that we form two layers of allele frequencies: the first layer is composed of independent draws from the BN distribution, and the second layer mixes these to create *p*_*i*1_, *p*_*i*2_, … , *p*_*iK*_ with coancestry **Λ**.

Let *S* be the number of components that will be mixed, ***W*** be the *S* × *K* matrix of mixture proportions, and **Γ** an *S* × *S* diagonal matrix. The entries of ***W*** are *w*_*su*_ where 0 ≤ *w*_*su*_ ≤ 1 and 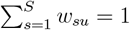 for *u* = 1, 2, … , *K*. The diagonal values of **Γ** are represented by *γ*_*s*_ where 0 ≤ *γ*_*s*_ ≤ 1, and all other values are 0. Suppose that for *i* = 1, … , *m* we generate

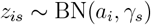

independently for *s* = 1, … , *S*, and we then set

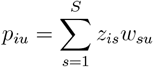

for *u* = 1, … , *K*. It can be verified that

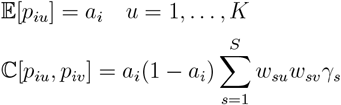

for *u, v* = 1, 2, … , *K*. By matching these equations with Eq. (6), one can see that if

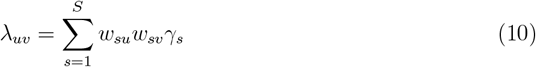

then *p*_*i*1_, *p*_*i*2_, … , *p*_*iK*_ has coancestry **Λ** as desired. In matrix terms, Eq. (10) is equivalent to

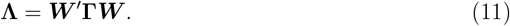

Therefore, the double-admixture method is based on the following optimization problem.

#### Problem 2

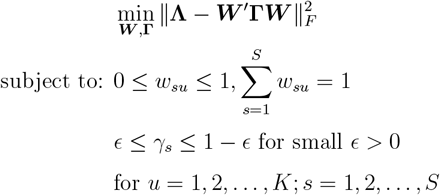

##### Algorithm 3

Calculating ***W*** and **Γ** in the double-admixture method

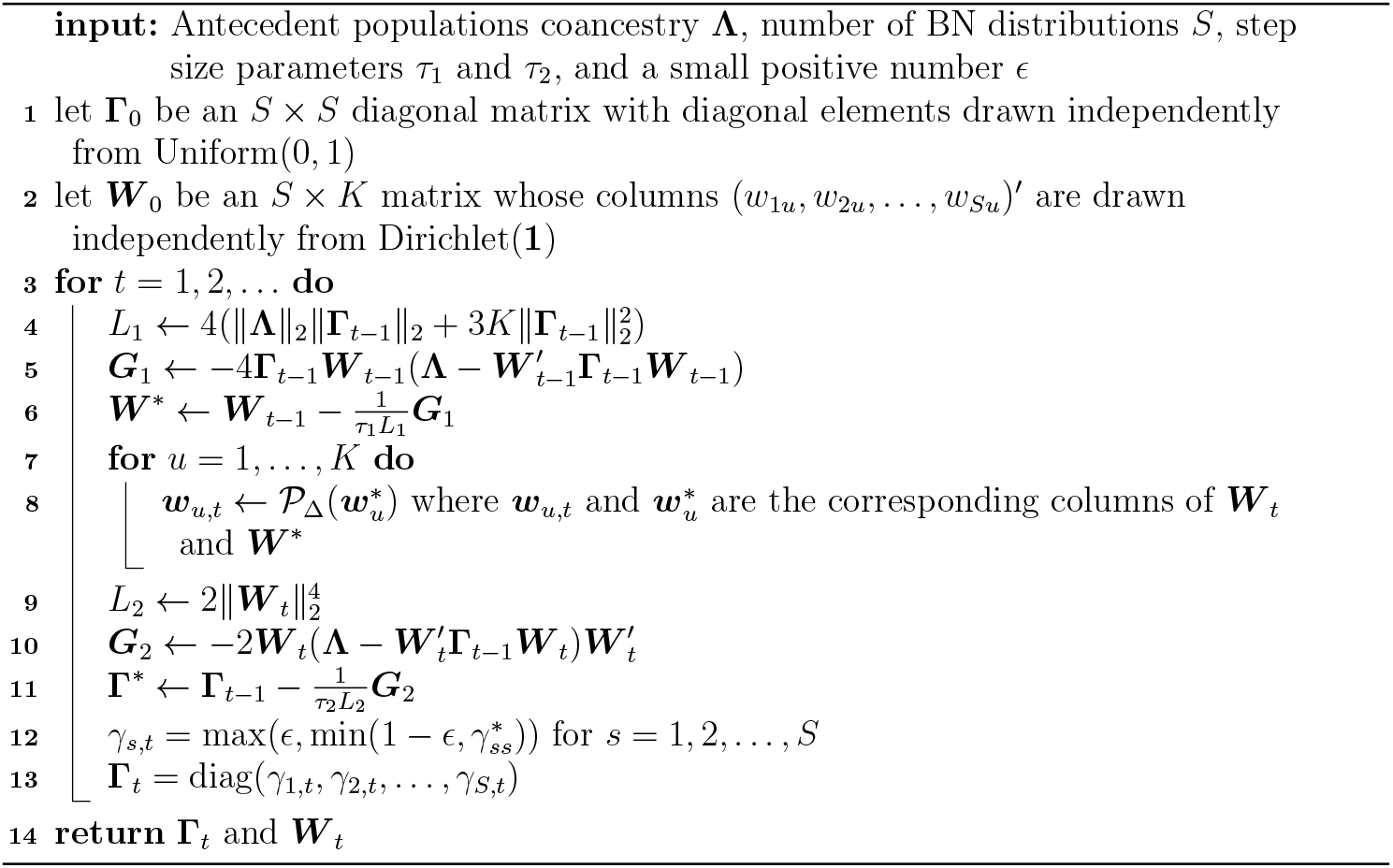

Here, we set *S* = 2*K*,*τ*_1_ = *τ*_2_ = 1.1, *ϵ* = 0.01; user should investigate their choices.

∥ · ∥_2_ denotes the spectral norm and 𝒫_Δ_ denotes projection onto the unit simplex (Appendix A.1).

We adapted the proximal alternating linearized minimization (PALM) method [26] to solve Problem 2, resulting in Algorithm 3 for calculating the parameters in the double-admixture method. Every sequence (***W*** _*t*_, **Γ**_*t*_)_*t*∈ℕ_ generated from Algorithm 3 is guaranteed to converge to a critical point. Integrating Algorithm 3 with the generative steps for *p*_*iu*_ described above, Algorithm 4 simulates antecedent population allele frequencies with the desired coancestry. In Appendix B.3, the PALM method is briefly introduced and the convergence of Algorithm 3 is proved.

##### Algorithm 4

The double-admixture algorithm for simulating ***P***

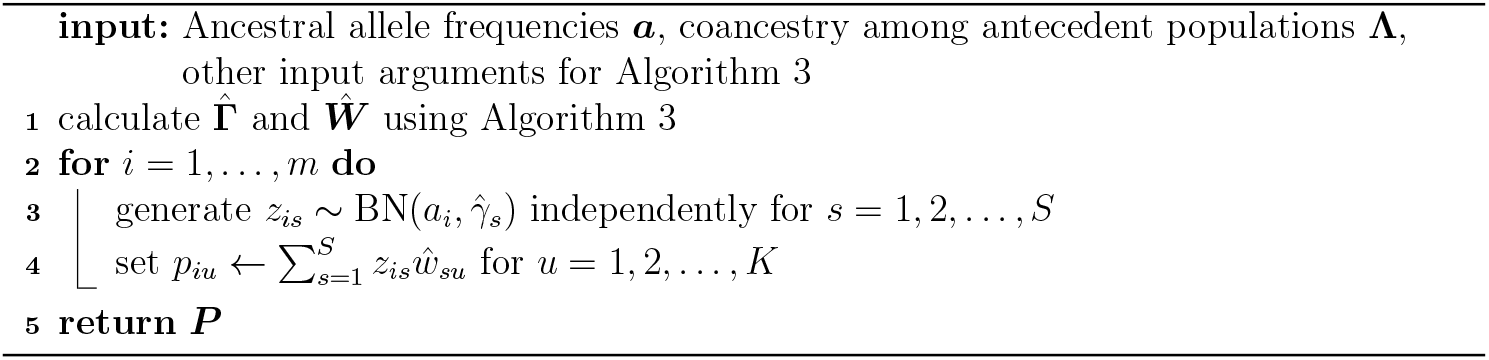

One possible drawback of the double-admixture method is that the approach relies on the existence of ***W*** and **Γ** so that **Λ** = ***W*** ′**Γ*W*** . We do not currently have a theoretical guarantee for such ***W*** and **Γ** (although one may exist since *S* can be made large). Therefore, we provide a complementary method in Appendix B.4, the NORmal To Anything (NORTA) approach [27], serving as a tool for simulating ***P*** when the double-admixture method is not applicable. It should be noted that the double-admixture method solves the optimization one time for the entire process so that its running time is independent of the number of loci *m*. In contrast, the NORTA method has to solve *K*× (*K*−1)*/*2 root-finding problems per locus and therefore has a complexity of 𝒪(*K*^2^*m*), rendering it significantly more time consuming. The performances of the double-admixture and NORTA methods are demonstrated in Appendix C.

Note that if we set **Γ** = **Λ** for a diagonal standard admixture **Λ** and ***W*** = ***I***_*K*_ (where ***I***_*K*_ is the *K* × *K* identity matrix), then the double-admixture method reduces to the BN sampling from Eq. (5), which produces valid antecedent population frequencies for the standard admixture model. From this observation, the double-admixture method can be viewed as a generalization of BN sampling.

### 2.5 Generating bootstrap datasets from realistic population structures

By utilizing the double-admixture method, we implemented the following algorithm to simulate genotypes from the super admixture model, shown in Algorithm 5. We assessed whether Algorithm 5 generates genotypes that satisfy the moment constraints imposed by the super admixture model in Appendix C. Algorithm 5 is especially useful when inputs ***a*, Λ**, and ***Q*** reflect real populations. When these parameters are unavailable one can utilize an admixture method to estimate ***Q*** and the method proposed here to estimate **Λ**. The ancestral allele frequencies can be estimated with simple sample means. We outline Algorithm 6, with the ALStructure algorithm for estimation of ***Q*** and the super admixture algorithm for estimation of **Λ**.

#### Algorithm 5

Generating genotypes ***X*** from the super admixture model

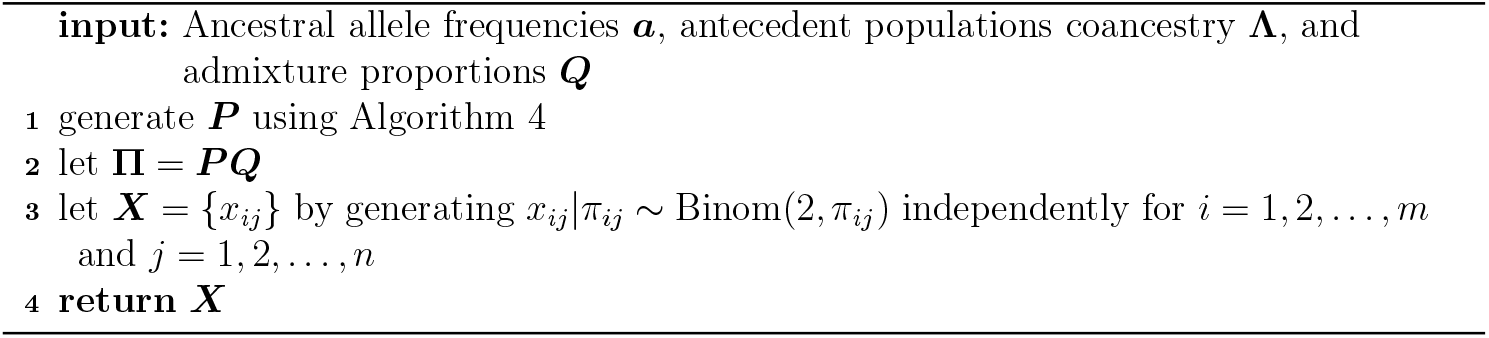

Line 1 can also be completed with the NORTA method, Algorithm B.4.

#### Algorithm 6

Generating bootstrap genotypes ***X***^*^ from observed genotypes ***X***

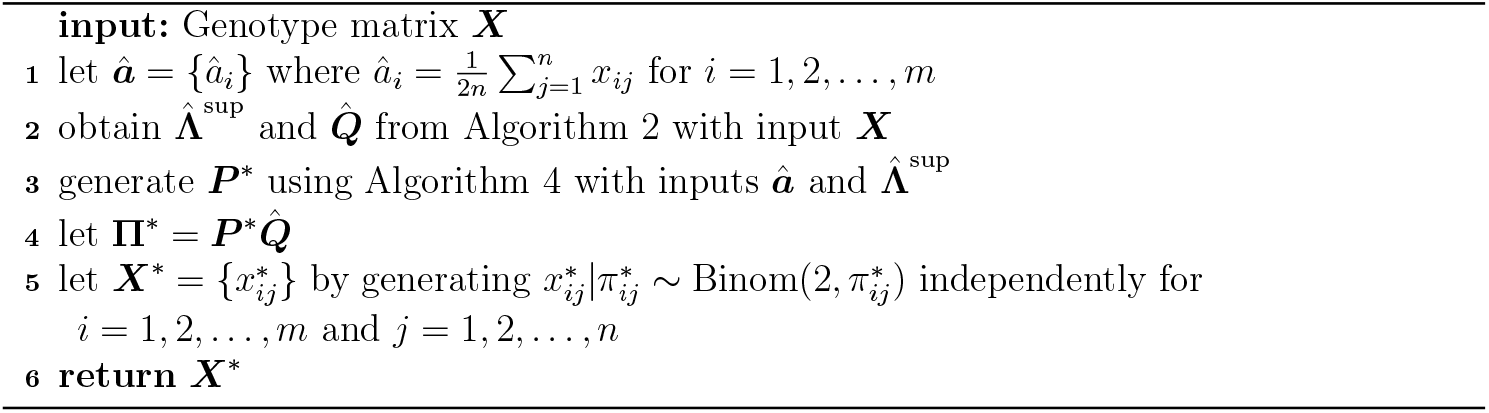

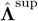 can be replaced with 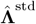 in Line 2, in which case the BN sampling from Eq. (5) is used in Line 3. Line 3 can also be completed with the NORTA method, Algorithm B.4, if using 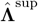.

We note that Algorithm 6 is a semi-parametric bootstrap simulation; Line 3 is semiparametric, **Π**^*^ is semi-parametric because 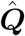 is nonparametric, and Line 5 is parametric. The output ***X***^*^ can be interpreted as a bootstrap replication of ***X***, where the population structure in ***X***^*^ recapitulates the structure in ***X***. The process that the bootstrap method recapitulates is not just resampled genotypes for a fixed matrix of estimated IAFs. Rather, the antecedent population allele frequencies are resampled, also leading to resampled IAFs, so both evolutionary and statistical resampling occur.

### 2.6 Significance test of coancestry among antecedent populations

Here, we develop a hypothesis test of the standard admixture model (null) versus the super admixture model (alternative). We show below that on real data sets the test results are highly significant against the null in favor of the alternative. In terms of model parameters, the test is defined as follows:

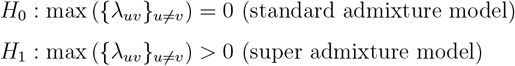

A straightforward test-statistic is 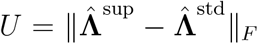. The larger *U* is, the more evidence there is against the null hypothesis in favor of the alternative hypothesis. In order to calculate a *p*-value for this test-statistic, we need to know the distribution of *U* when the null hypothesis is true. To this end, we adapt the bootstrap method of Algorithm 6, leading to Algorithm 7.

#### Algorithm 7

Hypothesis test of no coancestry among antecedent populations

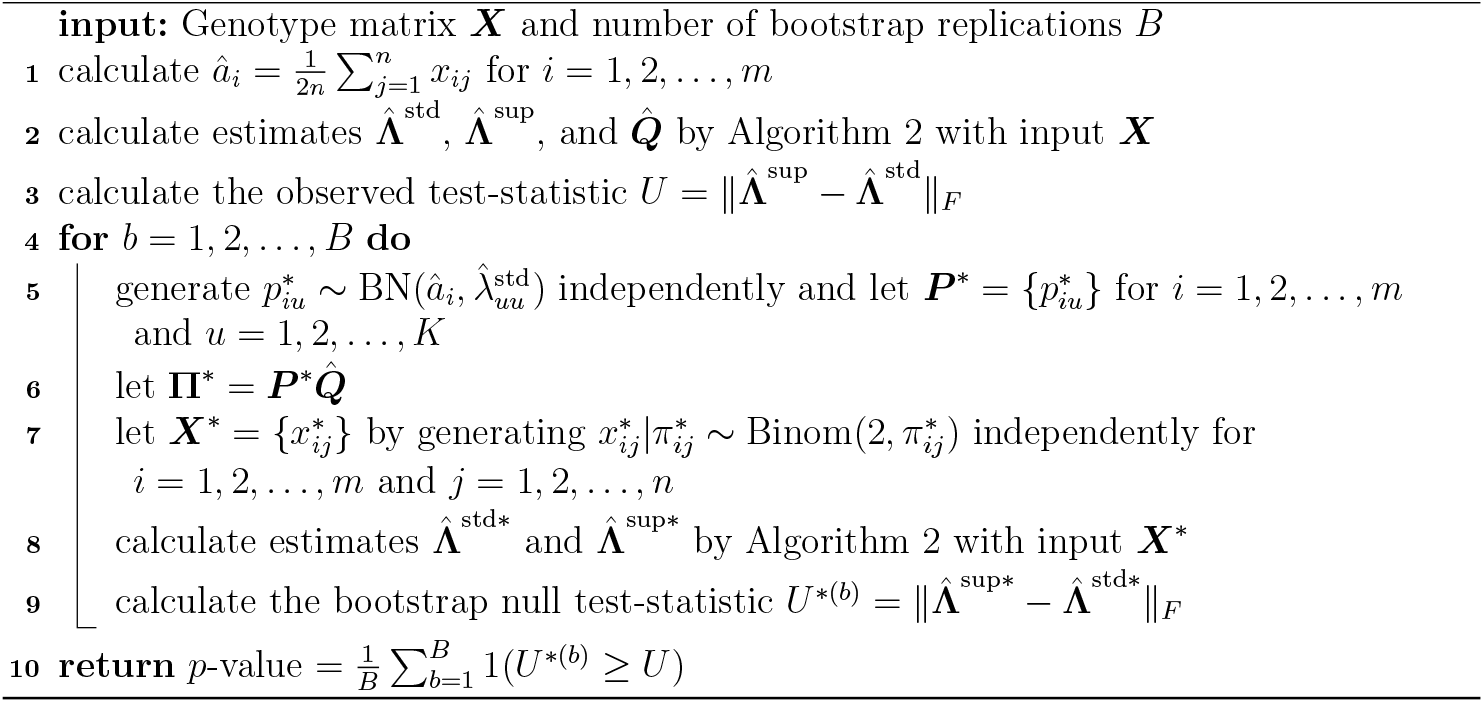

To evaluate the validity of the proposed test, we performed this hypothesis testing on various simulation designs (Appendix C). Our simulations show that the test produces valid *p*-values, which are conservative (Fig. C.4), meaning the test has a maximum type I error rate less than or equal to the nominal level of the test. On real data sets analyzed below, these *p*-values are small, so the conservative behavior that we observe in simulations does not appear to be relevant for populations with nontrivial levels of structure.

## 3 Analysis of human studies

We applied the super admixture framework to four published studies: the human genome diversity panel (HGDP) [19], the 1000 genomes project (TGP) [20], the Human Origins study (HO) [21], and a study on individuals with Indian ancestry (IND) [22]. Within the TGP study, we also analyzed a subset of admixed populations with American ancestry, denoted by AMR. While HGDP, TGP, and HO are sampled from ancestries throughout the world, the IND and AMR data sets are regionally sampled. This yielded five data sets that collectively represent a range of population structures and study designs. Discussions of the results on HO, AMR, and IND are in the main text, while HGDP and TGP are in Appendix D.

### 3.1 Calculations

We processed the data sets and performed quality control checks to produce a genotype matrix ***X*** for each as the starting point of our analysis (Appendix D.1). We next applied Algorithm 2 to ***X*** to obtain 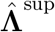 and 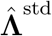 , the estimates of antecedent population coancestry for the super admixture and standard admixture models, respectively. We also calculated their corresponding individual-level coancestry estimates 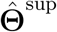 and 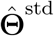 . As a part of Algorithm 2, we calculated the appropriate number of antecedent populations *K* using the structural Hardy-Weinberg method [12] (detailed in Appendix D.6). The values of *K* ranged from *K* = 11 for HO to *K* = 3 for AMR, which are consistent with earlier work [10, 12, 28]. Also, in Algorithm 2 we calculated estimates of the admixture proportion matrices 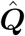 using the ALStructure method [10].

To evaluate the accuracy of 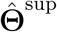 and 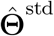 , we computed the OS estimate [4] of individual-level coancestry 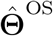 on each data set. The OS estimate of **Θ** is based on general assumptions and is a consistent estimator for arbitrary population structures under the appropriate conditions. Since 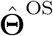 makes no assumptions about the distributions of the IAFs or coancestry parameters, it serves as a benchmark for our methods ^1^, allowing us to observe if the super admixture or standard admixture models lose information about individual-level coancestry relative to OS. As shown in Table D.1, the Frobenius-based distances from 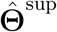 to 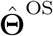 are about 10 to 40 times smaller than those from 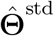 to 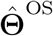. The distance from 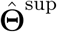 to 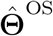 is smaller than is arguably practically relevant, meaning that 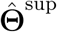 achieves the resolution of 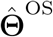 for practical purposes.

We carried out Algorithm 7 to perform a hypothesis test of the standard admixture model versus the super admixture model for all five datasets, with *B* = 1000 bootstrap iterations. For all data sets, no bootstrap null test-statistic was equal to or greater than the observed test-statistic, so *p*-value *<* 0.001 for all data sets. The bootstrap null test-statistics and observed test-statistic for all data sets are shown in Fig. D.9.

We applied Algorithm 6 to generate bootstrap replications ***X***^*^ from each data set’s genotype matrix ***X***. We applied the double-admixture method (Algorithm 4) and the NORTA method (Algorithm B.5) to include the performance of both methods. We computed the OS estimate 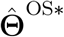 of individual-level coancestry for each ***X***^*^.

### 3.2 Visualizing results

We firstly visualized the results by making heatmaps of individual-level coancestry estimates 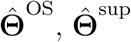, and 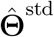. We also made heatmaps of 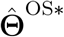 from bootstrap resampled genotypes using both the double-admixture and NORTA methods for generating antecedent population allele frequencies. These are displayed as follows: HO – Fig. 2, AMR – Fig. 4, IND – Fig. 6, HGDP – Fig. D.5, and TGP – Fig. D.7. It can be seen that for all data sets, 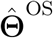 and 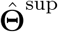 are qualitatively equivalent, which is quantitatively supported by Table D.1 showing they are very close. The estimates 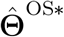 from the two bootstrap methods are also qualitatively equivalent to 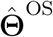 and 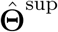. Finally, it can be seen that the standard admixture coancestry estimate 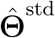 is not close to the other estimates, further indicating the standard admixture model is not sufficient for these data sets.

**Figure 4:**
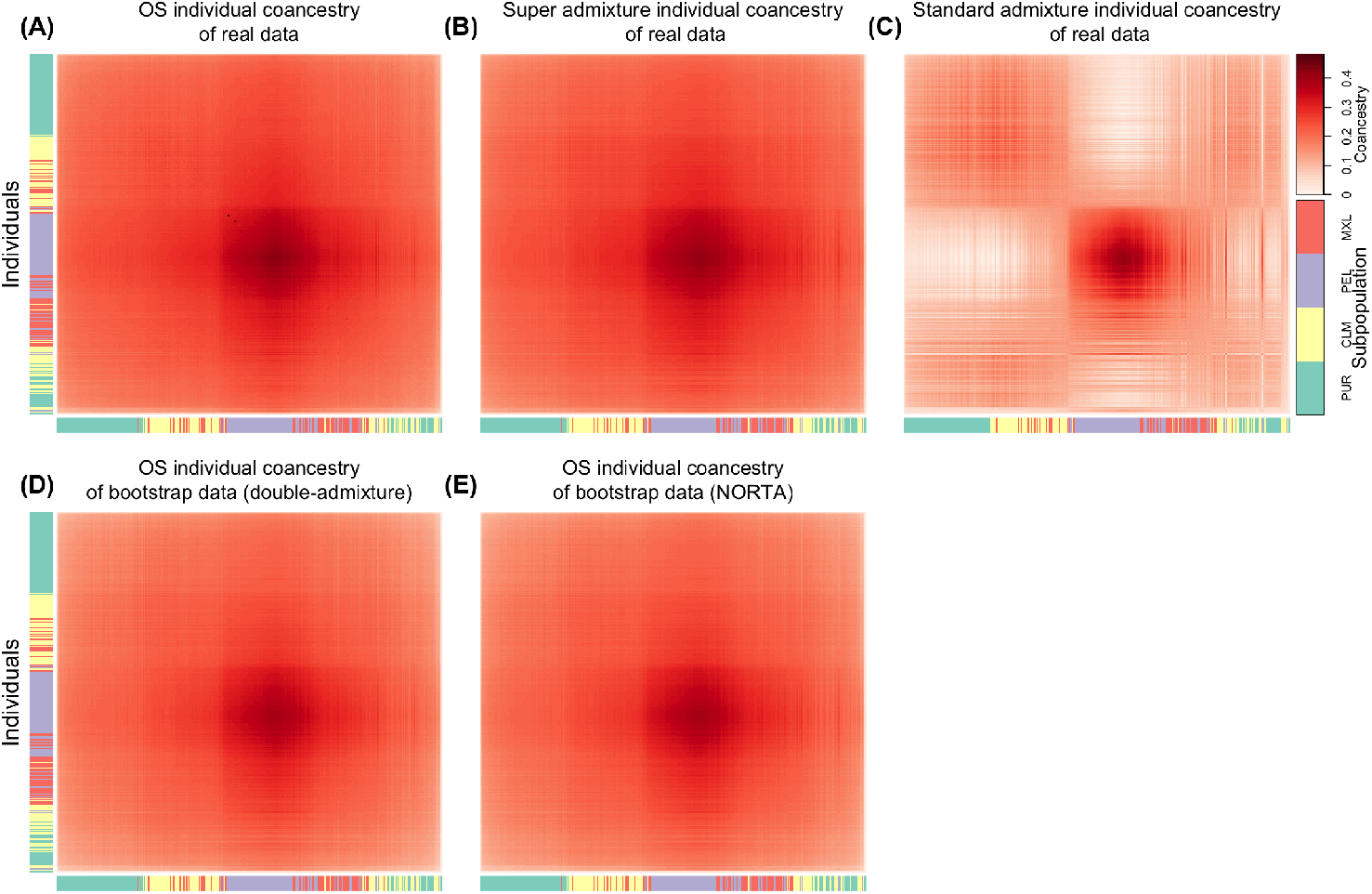
Heatmaps of individual-level coancestry estimates in the AMR data set.

We secondly visualized the results by building on the standard colored stacked bar plots of 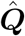 displaying the admixture proportions of the *K* antecedent populations for the individuals. In our case, we have additional information, which is the estimated antecedent population coancestry matrix 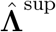 from the super admixture model. This matrix gives additional information about the relationship among the antecedent populations that we would like to visualize. The first way we visualized 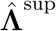 was create a heatmap of its values. We then constructed a dendogram built from 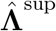 that is displayed above the stacked bar plot. This gives the user insight into the relatedness of the antecedent populations and connect them to the stacked bar plots. To this end, we calculated a distance matrix ***D*** from 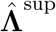 according to:

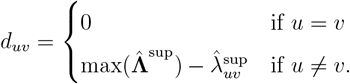

We then applied the standard agglomerative clustering method to ***D*** using “weighted pair group method with arithmetic mean” (WPGMA) to obtain a dendrogram. These are displayed in the data sets as follows: HO – Fig. 3, AMR – Fig. 5, IND – Fig. 7, HGDP – Fig. D.6, and TGP – Fig. D.8.

**Figure 5:**
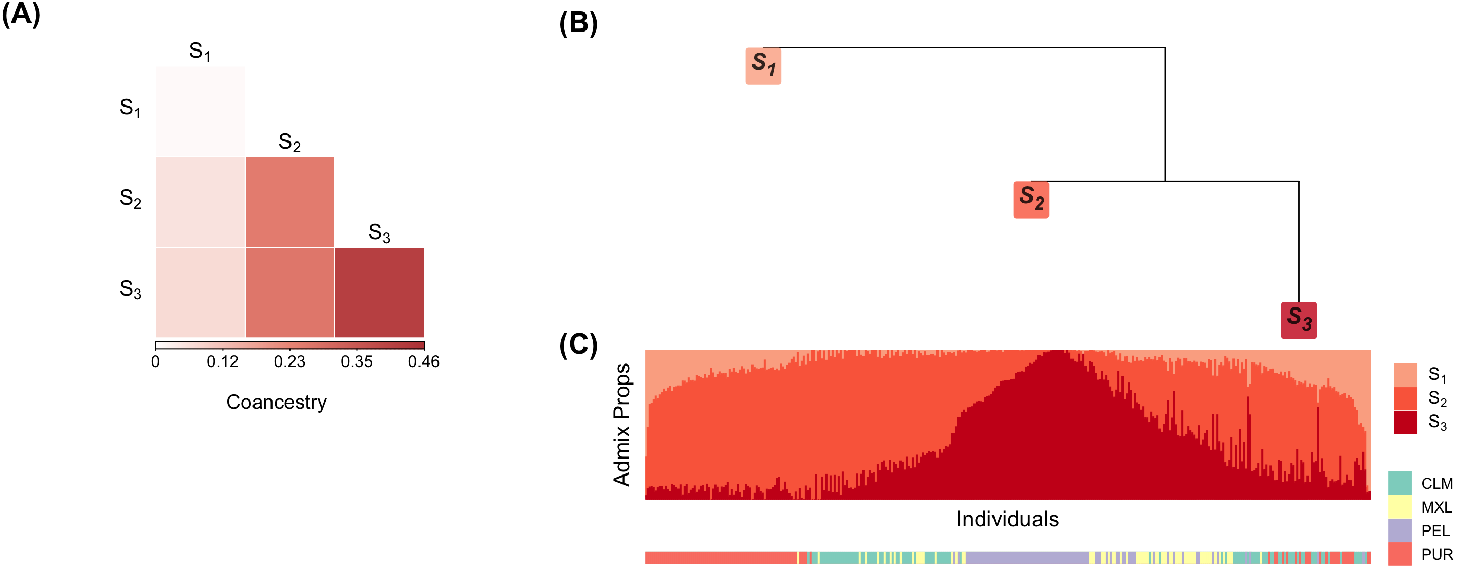
(A) Heatmap of antecedent population coancestry estimates in AMR. (B) Den-drogram representation of the antecedent population coancestry estimates. (C) Stacked bar plot of admixture proportions.

**Figure 6:**
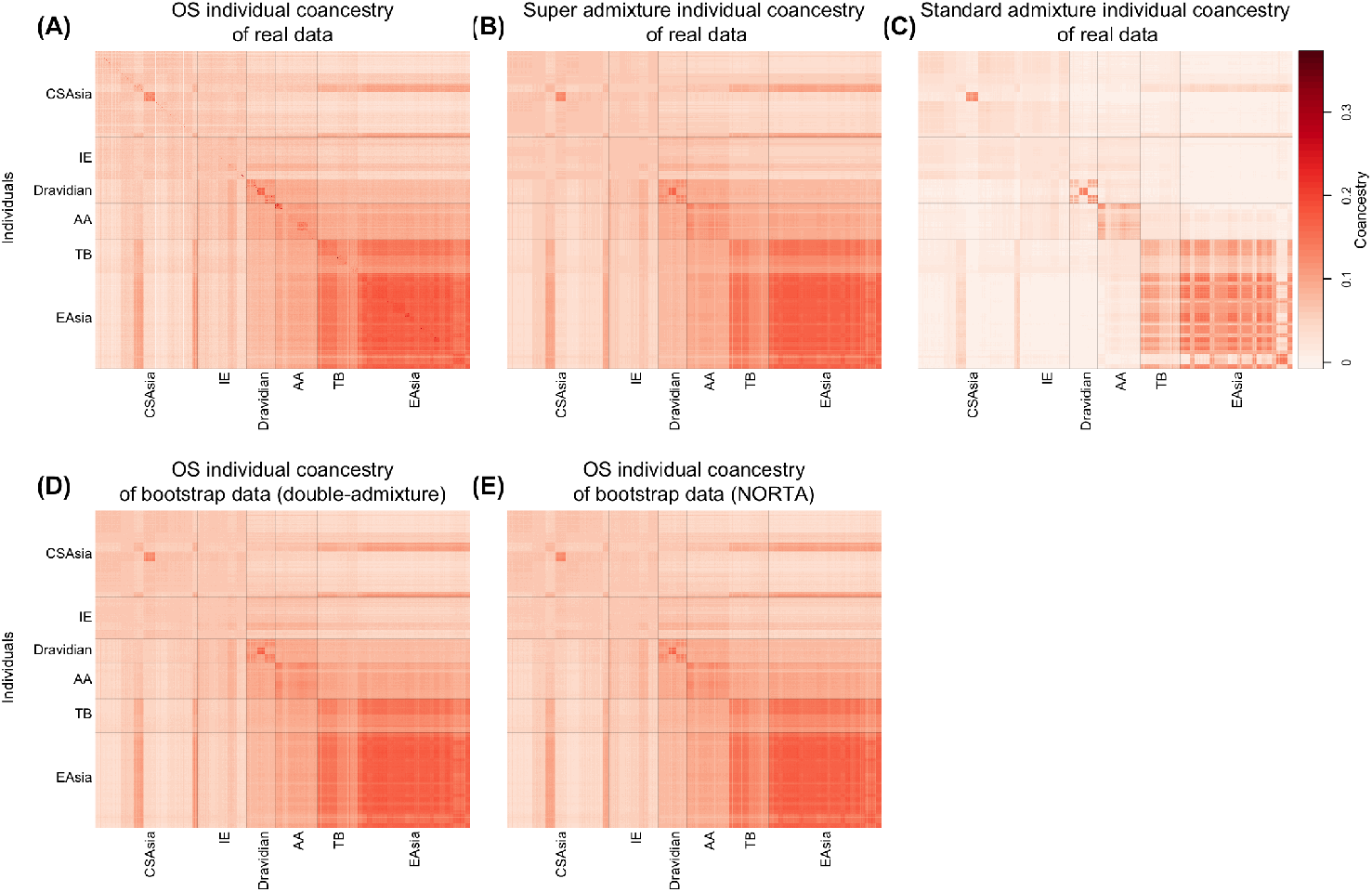
Heatmap of individual-level coancestry estimates in the merged data set of mainland Indians from IND, and Central/South Asians and East Asians from HGDP.

**Figure 7:**
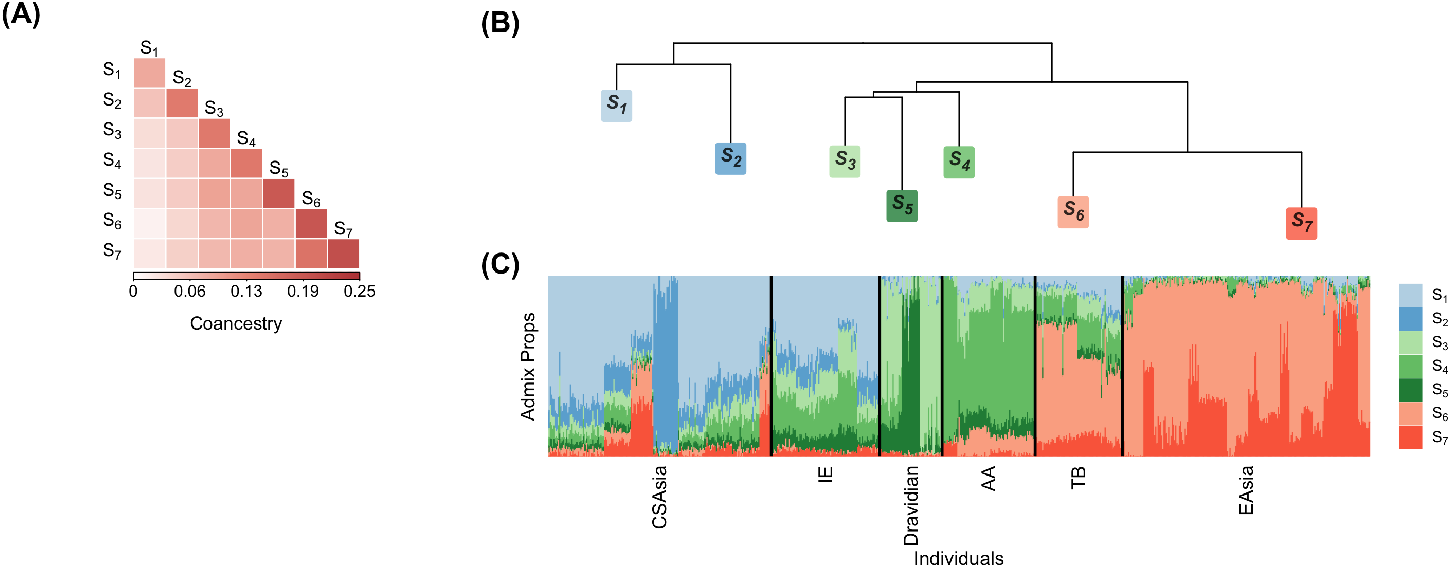
Heatmap of antecedent population coancestry estimates in the merged data set of mainland Indians from IND, and Central/South Asians and East Asians from HGDP. (B) Dendrogram representation of the antecedent population coancestry estimates. (C) Stacked bar plot of admixture proportions.

### 3.3 Human Origins (HO) study

The Human Origins datasets (HO) consists of 2124 individuals from 170 sub-subpopulations grouped into 11 subpopulations. We observed the estimated individual-level coancestry agrees with current knowledge of early human migrations [29–32]. In Fig. 2, we observed the first major split between Sub-Saharan Africa and North Africa. This split reflects the divergence between Sub-Saharan Africans and the rest of human populations resulting from an out-of-Africa migration around 50-60 kya. Another split occurred between South Asia and East Asia, revealing the separation between West Eurasians and East Asians around 40-45 kya. Among the East Asia clade, we identified that the Oceanians have highest coancestry within and lowest coancestry between other subpopulations, consistent with the theory that Oceanians split earliest from the rest of East Asians.

The coancestry among antecedent populations is also compatible with early human dispersals (Fig. 3). Specifically, in the dendrogram plot of the antecedent population coancestry (Fig. 3B), we note that the first branch split individuals from Sub-Saharan Africa represented by the antecedent populations *S*_1_ and *S*_2_ from individuals outside of Sub-Saharan Africa represented by the other antecedent populations. Individuals outside of Sub-Saharan Africa further branched off into two lineages: the West Eurasians represented by antecedent populations *S*_3_, *S*_4_ and *S*_5_, and the East Asians represented by antecedent populations *S*_6_ - *S*_11_. Then the Oceanians represented by the antecedent population *S*_9_ split off from the majority of East Asian ancestry, while the latter further diverged into present-day Asians (antecedent populations *S*_6_, *S*_7_, *S*_8_) and present-day Americans (antecedent populations *S*_10_ and *S*_11_).

### 3.4 Admixed individuals (AMR) from the 1000 Genomes Project (TGP)

The AMR subset of TGP has 353 individuals from four regions (Mexican-American (MXL): 65, Puerto Rican (PUR): 104, Colombian (CLM): 97, Peruvian (PEL): 87). The individual-level coancestry plot (Fig. 4) revealed that this dataset does not have a discrete population structure. Instead, the coancestry changes smoothly over individuals, indicating wide-ranging historical admixture events. This is consistent with the AMR population descending from European, Native American, and Sub-Saharan African ancestries during the post-Columbian era [33, 34].

In the analysis of the coancestry among antecedent populations (Fig. 5), we identified three major sources of ancestry: Sub-Saharan African ancestry represented by the antecedent population *S*_1_, West Eurasian ancestry represented by the antecedent population *S*_2_, and Native American ancestry represented by the antecedent population *S*_3_. The first split occurred between Sub-Saharan Africans (*S*_1_) and individuals outside of Sub-Saharan Africa (*S*_2_ and *S*_3_), and the second split between the West Eurasians (*S*_2_) and the Native Americans (*S*_3_). We also noted that the Puerto Ricans contain the highest amount of Sub-Saharan African ancestry; the Peruvians have the highest proportion of Native American ancestry; the Colombians and the Mexican-Americans display extensive variation in in their admixture proportions of European and Native American ancestry. Our observations were confirmed by previous analyses of AMR populations [28, 33, 34].

### 3.5 Indian (IND) study

We combined the mainland Indians from the IND study with the Central/South Asia and the East Asia populations from HGDP to study the relationship between present-day Indians and other populations in Asia. Our merged data set consists of 298 mainland Indians from fou linguistic groups (Indo-European (IE): 92, Dravidian: 53, Austro-Asiatic (AA): 79, Tibeto-Burman (TB): 74), together with 190 Central/South Asians and 210 East Asians from HGDP. Previous analyses of South Asian populations have shown that the Indo-European speakers show a considerable amount of the Western Eurasian relatedness and are ancestrally close to Central Asians. The Austro-Asiatic speakers and the Tibeton-Burman speakers were mixed from East Asian ancestry. The Tibeton-Burman speakers generally have significant genomic proportions derived from East Asian ancestry so that some Tibeton-Burman speakers can be difficult to distinguish from East Asian populations based on genome-wide measures of relatedness. Consistent with these findings [22, 35, 36], we observe a split between Indo-European speakers and the rest of mainland Indians in the heatmap of individual-level coancestry (Fig. 6). The Indo-European speakers and the Central/South Asians of HGDP have relatively similar levels of coancestry. The second split occurred between the Austro-Asiatic speakers and the Tibeto-Burman speakers. The Tibeto-Burman speakers and East Asians of HGDP have relatively similar levels of coancestry.

Our analysis reveals that there are three major branches of antecedent populations for this dataset (Fig. 7). The branch of antecedent populations *S*_1_ and *S*_2_ is most prevalent in Central/South Asians of HGDP and Indo-European speakers, suggesting this branch was at least partially derived from a West Eurasian source. The branch of the antecedent populations *S*_3_, *S*_4_ and *S*_5_ is widespread in Dravidian speakers and Austro-Asiatic speakers, indicating it is relevant to South Indian ancestry and Austro-Asiatic speaker ancestry. The third branch of the antecedent populations *S*_6_ and *S*_7_ likely represents East Asian ancestry due to its high prevalence in the Tibeto-Burman speakers and East Asians of HGDP.

## 4 Discussion

The super admixture framework is an extension of the highly used admixture model. It superposes coancestry among the admixed antecedent populations. It provides a forward generating probability process that encompasses random evolutionary, genealogical, and statistical sampling processes. The antecedent populations are modeled to have an arbitrarily complex coancestry. This allows the generation of individual-specific allele frequencies (IAFs) that capture complex population structures and permit the estimation of individual-level coancestry that is at the resolution of general individual-level coancestry and kinship estimators for arbitrarily complex structures.

There are numerous parameters estimated from genome-wide genotype data that relate to structure, such as coancestry, inbreeding, and *F*_ST_. When traits are included, one often estimates parameters in the context of genome-wide association studies [23, 37], genome-wide heritability [38–40] and polygenic risk scores [41, 42]. There does not exist a straightforward, general method for quantifying uncertainty among these various estimates. Within our framework, we have shown how to perform a bootstrap resampling method that randomly generates new genetic data that recapitulate population structure observed in real data. This bootstrap method may provide a way to formulate general methods for quantifying uncertainty in genome-wide genotype studies.

We developed a hypothesis test where one can test the standard versus super admixture model on real data. When we applied it to the five data sets analyzed here, all of them were highly significant in rejecting the standard admixture model in favor of the super admixture model. The individual-level coancestry estimates from the super admixture model also agreed with the general coancestry estimate, whereas the standard admixture individual-level coancestry estimates did not.

The stacked bar plot visualization of admixture proportions among individuals is ubiquitous in analyzing population structure. We showed here how the estimated antecedent population coancestry can be plotted with the stacked bar plot to visualize the relationship among the antecedent populations in conjunction with the bar plot. The admixture proportions among individuals are then interpretable in terms of the evolutionary history of the antecedent populations. We demonstrated this visualization on five data sets and showed how it agreed with known results on these human populations.

Understanding population structure in humans is one of the central problems in modern genetics. We demonstrated that the proposed super admixture framework is a powerful tool for learning admixed population coancestry, improving the analysis of genetic data from structured populations, bridging admixture with individual-level coancestry and kinship, and simulating new data reflecting a structured population. We anticipate that the super admixture framework will be useful in analyzing complex population structure in future applications.

## Resources

The *superadmixture* software package is available at https://github.com/StoreyLab/superadmixture. The results in this paper can be reproduced with code available at https://github.com/StoreyLab/superadmixture-manuscript-analysis.

## Acknowledgments

This work was supported in part by US National Institutes of Health grant R01 HG006448.

## Appendices

### A Supplementary theory

#### A.1 Mathematical notation and definitions

##### Definition 1

Given a vector ***x*** ∈ ℝ^*n*^, the 𝓁_*p*_ norm is defined as:

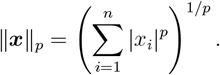

##### Definition 2

Given a matrix ***A*** ∈ ℝ^*m*× *n*^, the maximum singular value of ***A*** is denoted as *σ*_max_(***A***). The maximum and minimum eigenvalues of ***A*** are denoted as *λ*_max_(***A***) and *λ*_min_(***A***).

##### Definition 3

Given a matrix ***A*** ∈ ℝ^*m*× *n*^, the Frobenius norm of ***A*** is defined by:

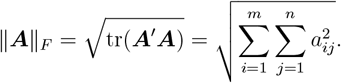

##### Definition 4

The induced matrix norm ∥***A***∥_*a*,*b*_ is defined as:

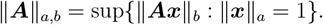

When *a* = *b* = 2, the induced matrix norm is the spectral norm:

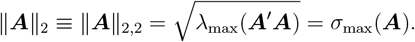

When *a* = *b* = 1, the induced matrix norm is the maximum absolute column sum of the matrix:

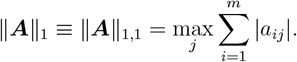

##### Definition 5

The proximal operator prox_f_ (***x***) is defined as

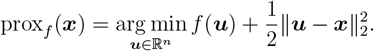

Let *f* (***x***) be an indicator function defined as

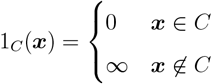

where *C* is a nonempty subset ℝ^*n*^. Let 𝒫_*C*_ denote the “projection onto *C* operator”. Then

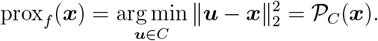

The *n*-dimensional unit simplex is defined as

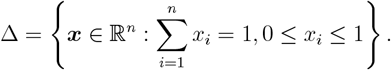

We define the projection onto the unit simplex operator 𝒫_Δ_ as

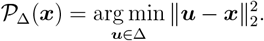

##### Definition 6

A subset *S* of ℝ^*n*^ is a real semi-algebraic set if there exists a finite number of real polynomial functions *g*_*ij*_, *h*_*ij*_: ℝ^*n*^ → ℝ such that

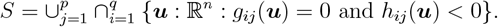

##### Definition 7

A function *f* : ℝ^*n*^ → (−∞, ∞] is called semi-algebraic if its graph

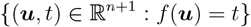

is a semi-algebraic subset of ℝ^n+1^.

#### A.2 Lemmas supporting the algorithms

##### Lemma 1

The definition of the induced matrix norm ∥ ***A***∥_*a*,*b*_ implies for any ***x*** ∈ ℝ^*n*^, the sub-additivity holds:

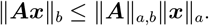

##### Lemma 2

Given matrices ***A, B*** ∈ ℝ^*m*× *n*^,

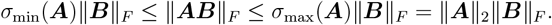

##### Lemma 3

Given a matrix ***A*** ∈ ℝ^*m*× *n*^,

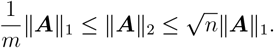

##### Lemma 4

Let *f* : ℝ^*n*^ → (−∞, ∞] be a proper and closed function. If *f* is semi-algebraic then it satisfies the Kurdyka-Lojasiewicz (KL) property at any point of dom(*f* ) [26].

### B Supplementary methods

#### B.1 Estimating individual-level pairwise coancestry

The OS estimate [4] begins with a measurement for allele matching between each pair of individuals:

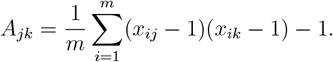

They prove that the expectation of *A*_*jk*_ is

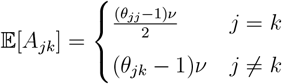

where 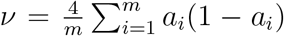. Let <u>*θ*</u> denote min_*j*,*k*_ *θ*_*jk*_. The Ochoa-Storey (OS) coancestry estimate utilizes <u>*θ*</u> = 0, which sets the reference population *T* to the most recent common ancestral (MRCA) population. When <u>*θ*</u> = 0, then *ν* = −<u>*A*</u> where <u>*A*</u> = min_*j*,*k*_ 𝔼[*A*_*jk*_]. Then OS estimator of coancestry is

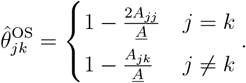

In general, −<u>*A*</u> = (<u>*θ*</u>− 1)*ν*. One can extend the OS estimator of coancestry for general values of <u>*θ*</u> as follows:

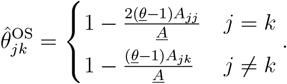

#### B.2 Estimating coancestry among antecedent populations

##### Proximal Forward-Backward (PFB) algorithm

Let *f* : ℝ^*n*^ → (−∞, +∞] be a be proper and closed function, let *h* : ℝ^*n*^ → (−∞, +∞) be convex and differentiable w ith a *L*-Lipschitz continuous gradient ∇*h*, i.e.,

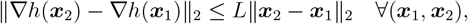

where *L* ∈ (0, ∞). Suppose that *f* (***x***) + *h*(***x***) → ∞ as ∥***x***∥_2_ → ∞. The problem is to identify:

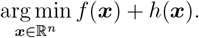

It has been shown that this problem can be solved by the PFB algorithm [24]. Every sequence (***x***_*t*_)_*t*∈ℕ_ generated by the following constant-step forward-backward algorithm converges to a solution to the problem.

###### Algorithm B.1

The constant-step forward-backward algorithm

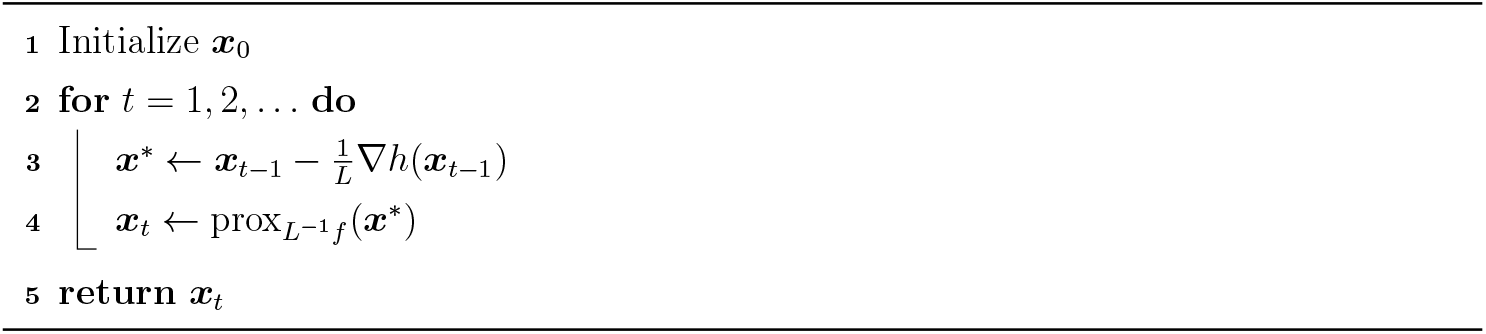

prox(·) denotes the proximal operator (Appendix A.1).

##### Solving Problem 1 by PFB

Problem 1 is equivalent to the problem of finding the minimizer of *f* (**Λ**) + *h*(**Λ**) where

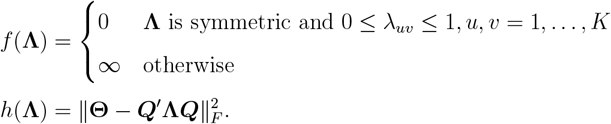

The function *f* is proper and closed because dom(*f* ) is nonempty and closed. The function *h* is differentiable w ith a c ontinuous g radient ∇*h* = −2***Q***(**Θ** − ***Q*** ′**Λ*Q***)***Q***′. ∇ *h* i s Lipschitz continuous with 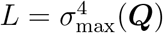.

*Proof*. We note that

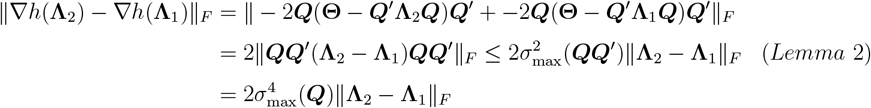

Therefore, we can employ the PFB algorithm to solve Problem 1. The proximal operator 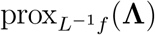 can be calculated as 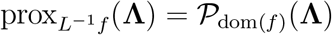. This implies

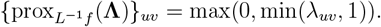

This leads to Algorithm 1, where we have now proved that every sequence (**Λ**_*t*_)_*t*∈ℕ_ converges to a solution.

##### Solving Problem B.1 by PFB

We can formulate the estimation of coancestry among antecedent populations under the standard admixture model as follows.

###### Problem B.1

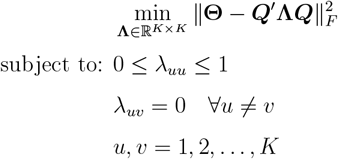

It is straightforward to see that Problem B.1 is identical to identifying the minimizer of *f* (**Λ**) + *h*(**Λ**) where

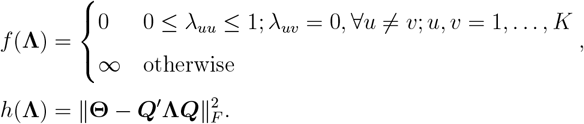

The function *f* is proper and closed and *h* is differentiable with a continuous gradient ∇*h* = −2***Q***(**Θ** − ***Q***′**Λ*Q***)***Q***′. The gradient ∇*h* is Lipschitz continuous with 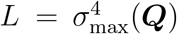. By Appendix A.1, 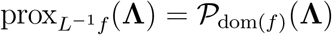. This implies:

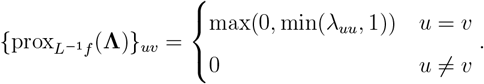

This leads to Algorithm B.2, where every sequence (**Λ**_*t*_)_*t*∈ℕ_ converges to a solution.

###### Algorithm B.2

Estimating **Λ** for the standard admixture model given **Θ** and ***Q***

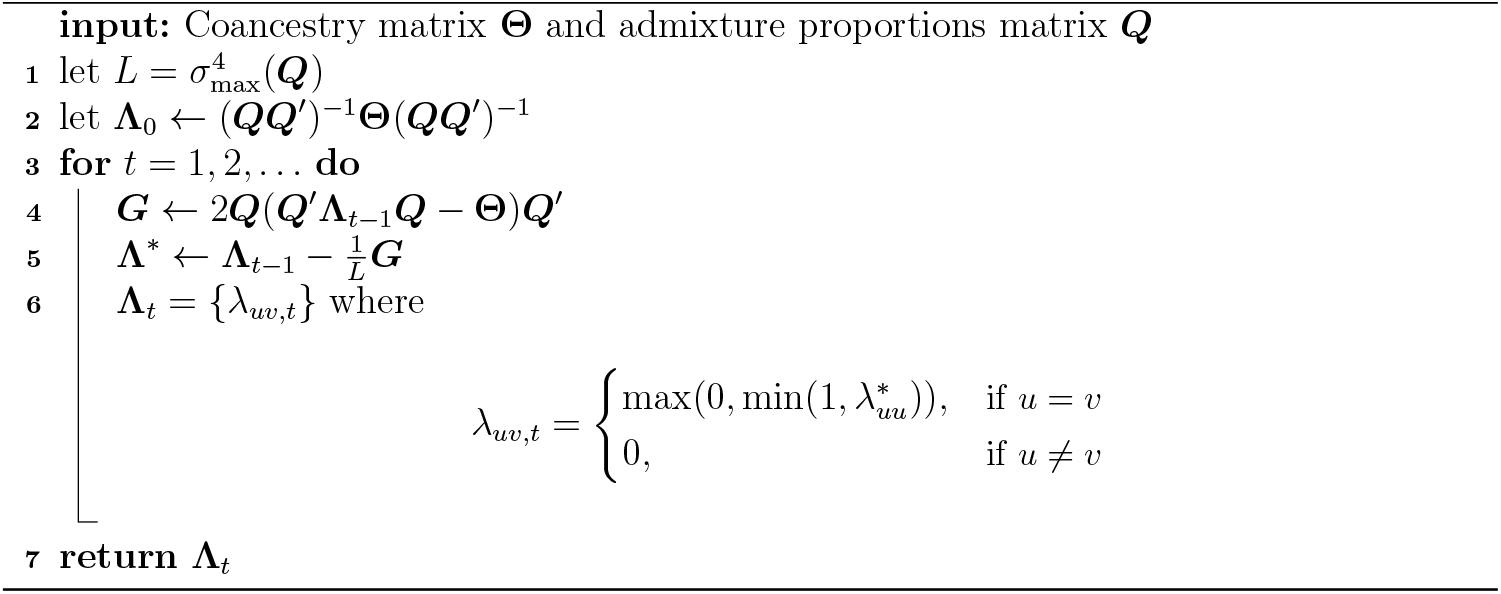

*σ*_max_(·) denotes the maximum singular value (Appendix A.1).

#### B.3 Estimating Parameters in the Double-Admixture Model

##### Proximal Alternating Linearized Minimization (PALM) aglorithm

Let *f* : ℝ^*n*^ → (−∞, +∞] and *g* : ℝ^*m*^ → (−∞, +∞) be closed functions. Let *h* : ℝ^*n*^ × ℝ^*m*^ → ℝ be a continuously differentiable function. The problem is to find a solution to:

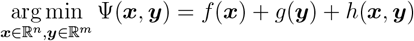

over all (***x, y***) ∈ ℝ^*n*^ × ℝ^*m*^. It has been shown that this problem can be solved by the Proximal Alternating Linearized Minimization (PALM) algorithm. Assume that:

i. 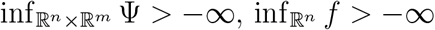, and 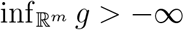.
ii. Ψ is a Kurdyka-Lojasiewicz function (see Appendix A.1).
iii. *h* is twice continuously differentiable.
iv. There exists convex and compact sets 𝒞_***x***_ and 𝒞_***y***_ such that ***x***_*t*_ ∈ 𝒞_***x***_ and ***y***_t_ ∈ 𝒞_***y***_ for all *t* ∈ ℕ.
v. For any fixed ***y*** the partial gradient ∇_***x***_*h*(***x, y***) is Lipschitz continuous with moduli *L*_1_(***y***) over the domain 𝒞_***x***_, that is

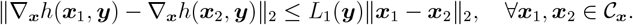 Likewise, for any fixed ***x*** the partial gradient ∇_***y***_*h*(***x, y***) is Lipschitz continuous with moduli *L*_2_(***x***) over the domain 𝒞_***y***_.
vi. For *i* = 1, 2 there exists 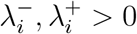 such that:

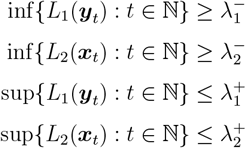

We note that these assumptions are not exactly the same as the assumptions specified in ref. [26]. We modified the original assumptions to align the PALM algorithm to our setting. Following the proof provided in ref. [26], one can show that if these assumptions are met, the sequence (***x***_*t*_, ***y***_*t*_)_*t*∈ℕ_ generated by Algorithm B.3 converges to a critical point of Ψ.

###### Algorithm B.3

The general PALM algorithm

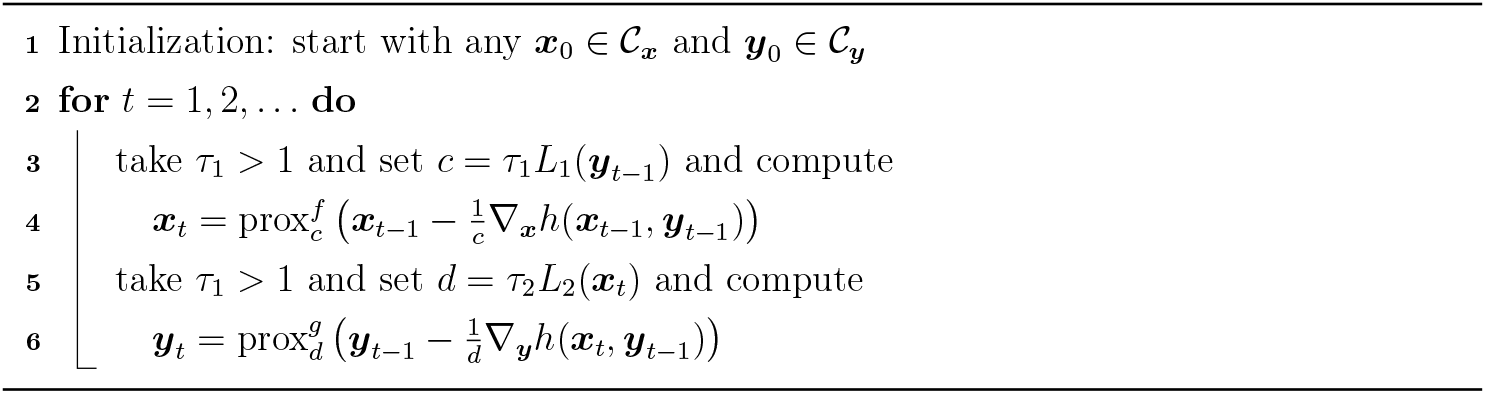

prox(·) denotes the proximal operator (Appendix A.1).

##### Solving Problem 2 by PALM

Problem 2 is identical to identifying the minimizer of *f* (***W*** ) + *g*(**Γ**) + *h*(***W*** , **Γ**) where

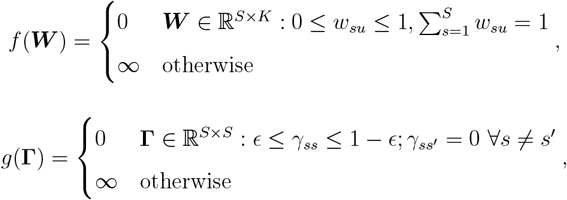

and *h* is defined as 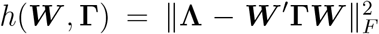 . Define 𝒞_***W***_ = {***W*** ∈ ℝ^*S*× *K*^ : *w*_*su*_ ≥ 0,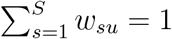}. Define 𝒞_**Γ**_ = {**Γ** ∈ ℝ^*S*× *S*^ : *ϵ* ≤ *γ*_*ss*_ ≤ 1 − *ϵ*; *γ*_*ss*_*′* = 0 ∀*s s*′}. We note that both functions ***W*** → ∇_***W***_ *h*(***W*** , **Γ**) and **Γ** → ∇_**Γ**_*h*(***W*** , **Γ**) are continuous. Indeed,

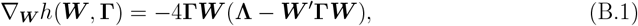

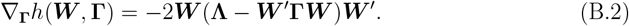

For all ***W*** _1_, ***W*** _2_ ∈ *C*_***W***_ ,

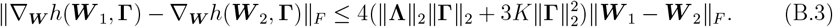

For all **Γ**_1_, **Γ**_2_ ∈ *C*_**Γ**_,

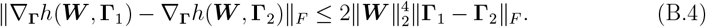

Eqs. (B.3) and (B.4) are proved in the following paragraphs. By Appendix A.1, 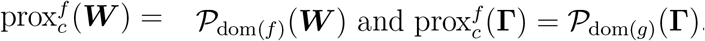. This implies

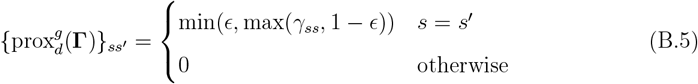

and

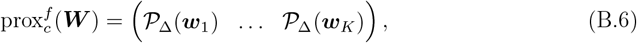

where ***w***_1_, … , ***w***_*K*_ are columns of ***W*** . Applying Eqs. (B.1) to (B.6) to Algorithm B.3, we arrive at Algorithm 3 for solving Problem 2.

##### Proving the convergence of Algorithm 3

To prove the convergence, we need to show all assumptions of PALM hold. It is obvious that the assumptions (i), (iii) and (iv) hold.

###### Proof of assumption (ii)

Ψ is a KL function. By Lemma 4, we note that the objective function *H* is a real polynomial function, hence semi-algebraic. For the indicator function *f* , we observe that the domain of *f* is defined by 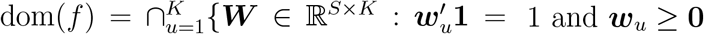. Hence, dom(*f* ) is a semi-algebraic set, so *f* is a semi-algebraic function. For the indicator function *g*, we observe that the domain of *g* is defined by 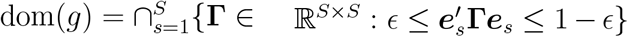, where ***e***_1_ = (1, 0, … , 0), …, ***e***_*S*_ = (0, 0, … , 1). Hence, dom(*g*) is a semi-algebraic set, so *g* is a semi-algebraic function. Thus, Ψ = *f* +*g* +*H* is a semi-algebraic function, and Ψ satisfies the KL property of any point of its domain.

*Proof of assumption (v)*: To prove the assumption (v), we are to show Eqs. (B.3) and (B.4). We note that for all ***W*** ∈ 𝒞_***W***_ , by the definition of the induced matrix norm and Lemma 3, we have 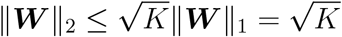. For all ***W*** _1_, ***W*** _2_ ∈ 𝒞_***W***_ :

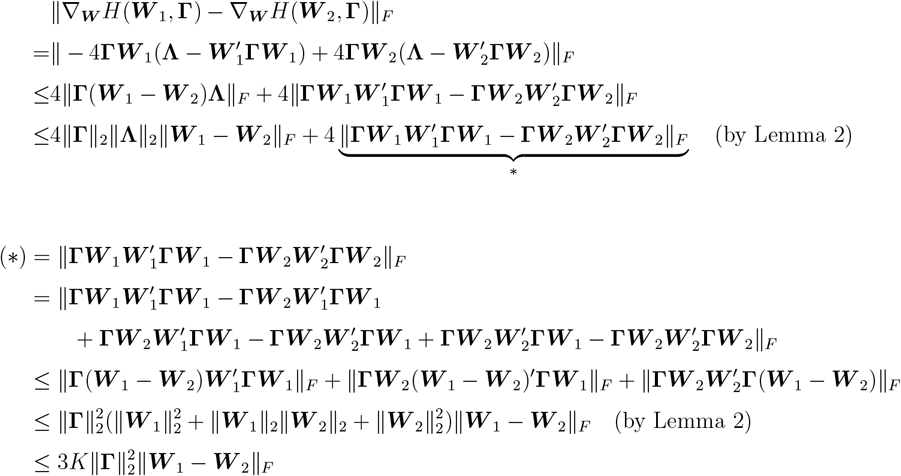

Therefore,

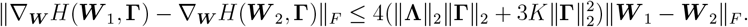

For all **Γ**_1_, **Γ**_2_ ∈ ℝ^*K*× *K*^ ,

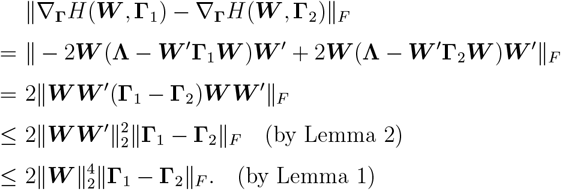

###### Proof of assumption (vi)

Since ***W*** _*t*_ ∈ 𝒞_***W***_ and **Γ**_*t*_ ∈ 𝒞_**Γ**_ for all *t* ∈ ℕ, and 𝒞_***W***_ and 𝒞_**Γ**_ are compact sets, 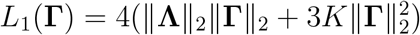 and 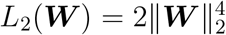 are bounded. By Lemma 3, 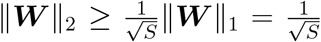 and 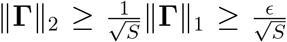 . Therefore, 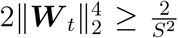 and 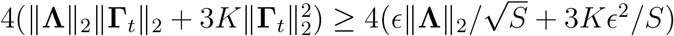 for all *t* ∈ ℕ.

#### B.4 Simulating antecedent population coancestry through NORmal To Anything (NORTA)

##### NORmal To Anything (NORTA) algorithm

The NORmal To Anything (NORTA) algorithm is a transformation-based method for generating random vectors with given marginal distributions and a given covariance matrix. The goal of the NORTA method is to define a *K*-dimensional random vector ***X*** with the following properties:

i. *X*_*u*_ ∼ *F*_*u*_, *u* = 1, … , *K*, where {*F*_*u*_} are marginal cumulative distribution functions,
ii. ℂ(***X***) = **Σ**_***X***_ = {*σ*_*uv*,***X***_}.

The NORTA algorithm represents ***X*** as a transformation of a *K*-dimensional, standard multivariate normal vector ***Z*** = (*Z*_1_, *Z*_2_, … , *Z*_*K*_)′ with covariance matrix ℂ(***Z***) = **Σ**_***Z***_ = {*σ*_*uv*,***Z***_}.

###### Algorithm B.4

NORTA algorithm

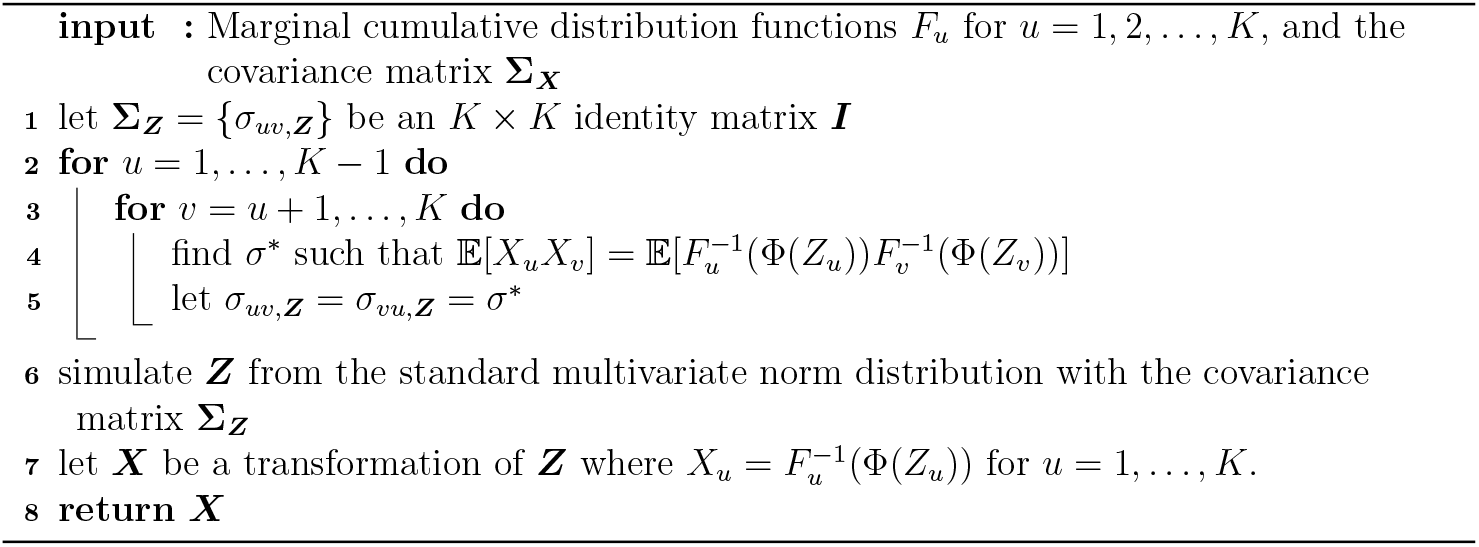

Φ is the univariate Normal(0, 1) cumulative distribution function (cdf); 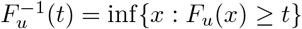 denotes the inverse cdf.

Note that 𝔼[*X*_*u*_], 𝔼[*X*_*v*_], 𝕍(*X*_*u*_) and 𝕍(*X*_*v*_) are determined by *F*_*u*_ and *F*_*v*_, implying 𝔼[*X*_*u*_*X*_*v*_] is determined by *F*_*u*_ and *F*_*v*_. Let *Φ*_*σ*_ denote the standard bivariate normal probability density distribution with the correlation (and also covariance in this case) *σ*. Then

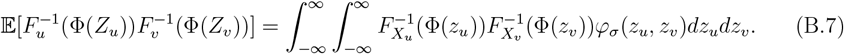

Determining *σ* to yield the desired covariance is equivalent to solving the root of the function

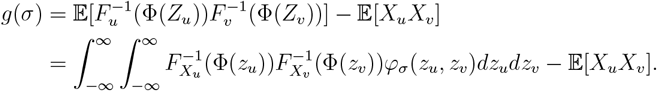

We will denote this root by *σ*^*^ where *g*(*σ*^*^) = 0.

##### Applying NORTA to generate coancestry among antecedent populations

We applied the NORTA algorithm with ***X*** = ***p***_i_, *F*_*u*_ = BN(*a*_*i*_, *λ*_*uu*_), and *σ*_*uv*,***X***_ ≡ *a*_*i*_(1 − *a*_*i*_)*λ*_*uv*_ for *i* ∈ 1, … , *m*. In this scenario,

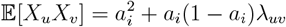

We note that there is no closed form solution for *σ*^*^. We adopted the Newton method to perform a numerical search for *σ*^*^. Let

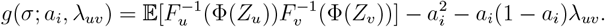

It follows that

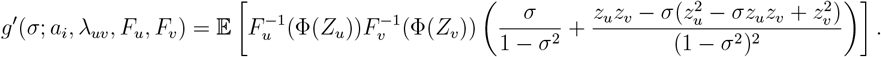

The Newton iteration for finding *σ*^*^ is then given by

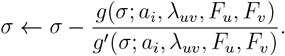

We calculated *g*(*σ*; *a*_*i*_, *λ*_*uv*_, *F*_*u*_, *F*_*v*_) and *g*′(*σ*; *a*_*i*_, *λ*_*uv*_, *F*_*u*_, *F*_*v*_) via numeric integration, leading to Algorithm B.5 for simulating antecedent population allele frequencies with the desired coancestry.

### C Supplementary numerical results

#### C.1 Generating Λ

To simulate **Λ** under the super admixture model, we simulated a *K* × *K* matrix ***A*** with elements drawn independently from Uniform(0, 0.3). We then set **Λ** = ***A***′***A***. To simulate **Λ**

##### Algorithm B.5

NORTA algorithm for simulating ***P***

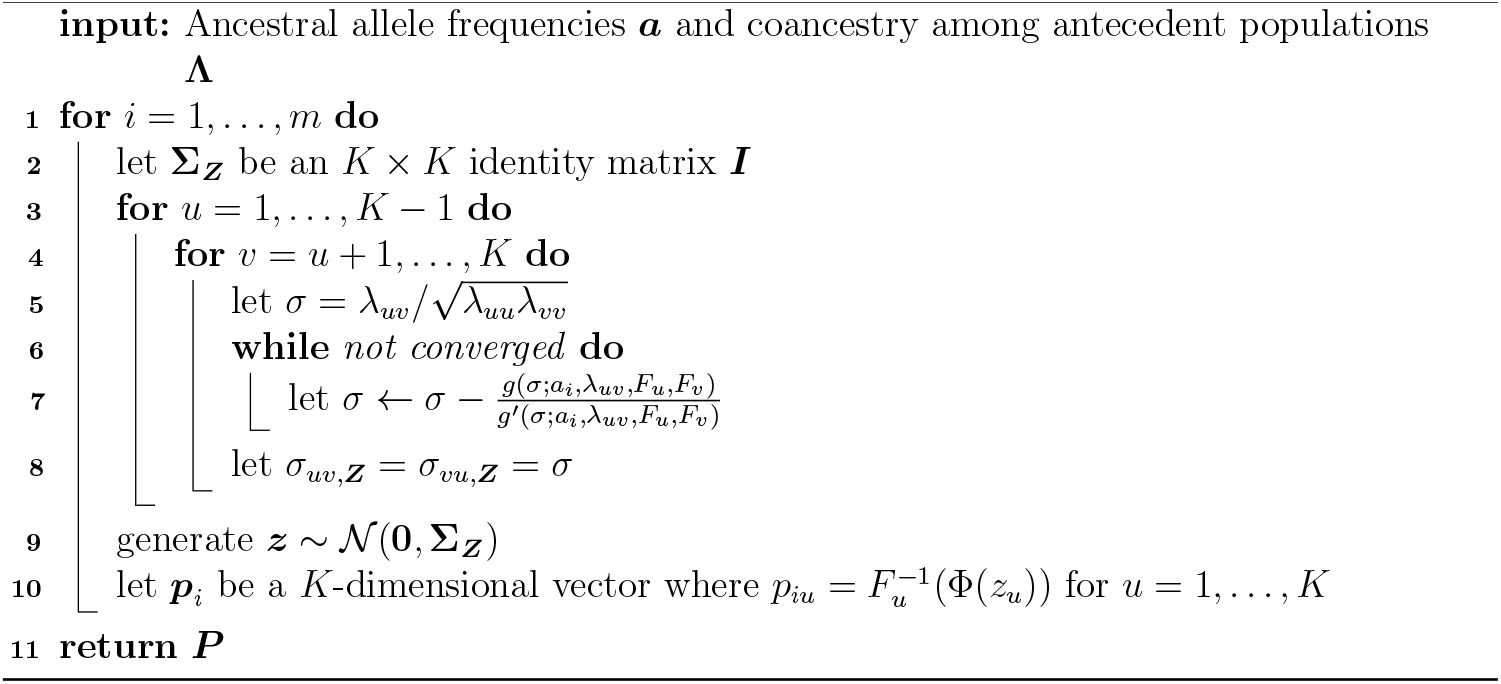

Φ is the univariate Normal(0, 1) cumulative distribution function (cdf); 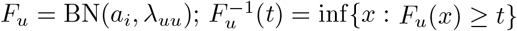 denotes the inverse cdf of *F*_*u*_.

under the standard admixture model, we let **Λ** be a diagonal matrix whose diagonal elements are generated independently from Uniform(0, 1). For both scenarios, we varied *K* = 3, 6, 9 and sampled 100 instances of **Λ** for each *K*. Our simulation resulted in 300 instances of **Λ** under the standard admixture model and 300 instances of **Λ** under the super admixture model.

#### C.2 Generating *Q*

We adopted the spatial model from ref. [4] to generate admixture proportions reflecting real data. This model represents the admixture process as a diffusion on a one-dimensional geography. It assumes *K* independent populations equally spaced at positions *x*_0_, *x*_0_ + 1, … , *x*_0_ + *K* − 1 on an infinite line. If all populations begin to diffuse at time *t* = 0 at the same diffusion rate, then population *u* will be distributed as a Gaussian with mean *µ*_*u*_ = *x*_0_ + *u* − 1 and standard deviation *σ*. Therefore, under the spatial model an individual *j* sampled at the position *j* will have the admixture proportions shown as follows:

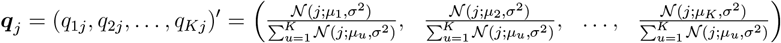

where 𝒩 (; *µ, σ*^2^) denotes a Normal(*µ, σ*^2^) distribution. We chose *σ*^2^ = 0.5 and *n* = 2000 in our simulations.

#### C.3 Evaluating algorithms for estimating coancestry among antecedent populations

To evaluate Algorithms 1 and B.2, we generated 300 unique combinations of (**Λ, *Q***) under the standard admixture model and 300 unique combinations of (**Λ, *Q***) under the super admixture model. For each pair of (**Λ, *Q***), we calculated the corresponding **Θ** = ***Q***′**Λ*Q***. If the super admixture model is assumed, we applied Algorithm 1 with a random initial matrix **Λ**_0_ to estimate **Λ**. If the standard admixture model is assumed, we applied Algorithm B.2 with a random initial matrix **Λ**_0_ to estimate **Λ**. We recorded values of **Λ**_*t*_ per iteration. We quantified the differences between **Λ** and **Λ**_*t*_ by 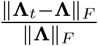 , and visualized the change over iterations in Fig. C.1. We validated that both algorithms are capable of generating a sequence of **Λ**_*t*_ such that the difference between **Λ** and **Λ**_*t*_ decreases as *t* → ∞.

**Figure C.1:**
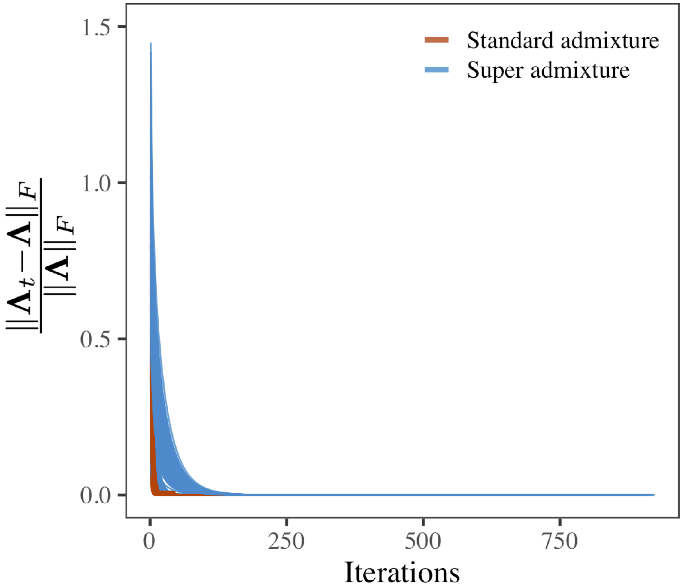
The convergence of Algorithms 1 and B.2.

#### C.4 Evaluating algorithms for antecedent population allele frequencies

To evaluate Algorithms 4 and B.5,, we assessed whether they could generate allele frequencies that satisfy the moments of the super admixture model:

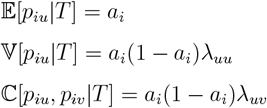

To achieve this, we generated 300 unique combinations of (*a*, **Λ**) where *a* is a scalar and is simulated from Uniform(0, 1); **Λ** assumes the super admixture model and is simulated as previously described. For each pair of (*a*, **Λ**), we generated *B* = 100, 000 replications of the *n*-vector allele frequenecies ***p***^(*b*)^ from the double-admixture method (Algorithm 4) or from the NORTA method (Algorithm B.5). Then we calculated the empirical mean and the empirical covariance matrix as 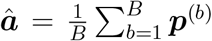 and 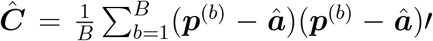, respectively. We measured the differences between empirical moments and the desired moments by 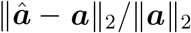 and 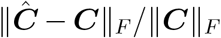 , where ***a*** denotes a *K* × 1 dimensional vector whose entries are all equal to *a* and ***C*** = *a*(1 − *a*)**Λ**. We found that 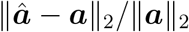 is generally less than 0.02 and 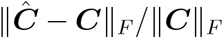 is generally less than 0.04 for both algorithms (Fig. C.2). These findings confirmed the performance of both algorithms.

**Figure C.2:**
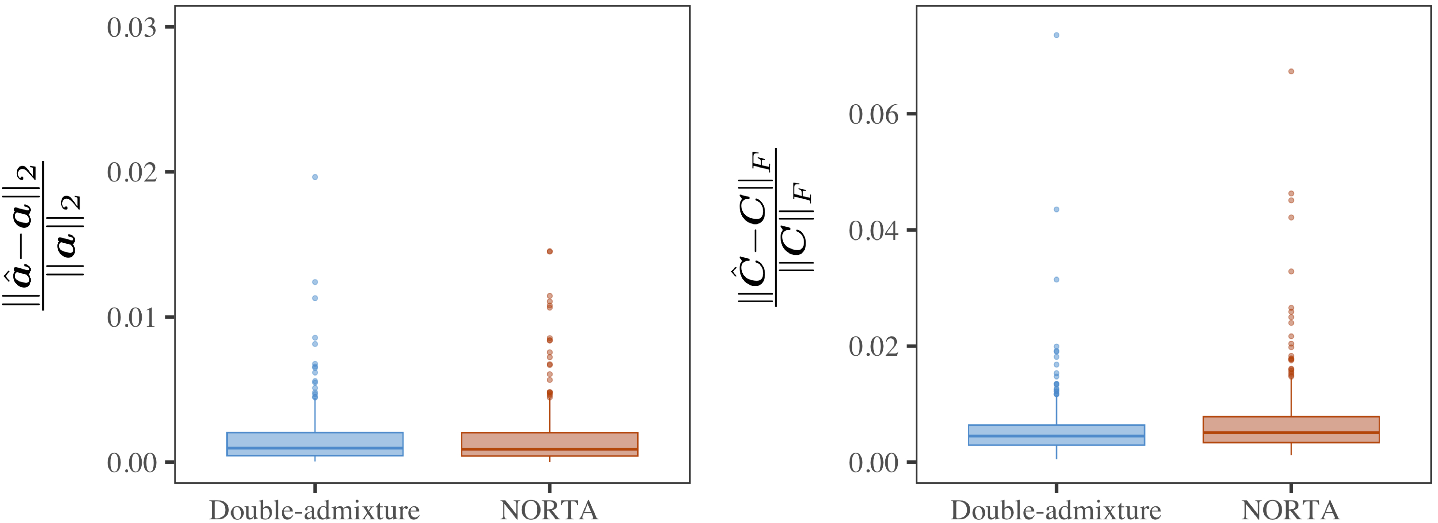
Box-plots of 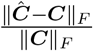 and 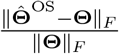 across 300 simulations.

#### C.5 Evaluating the algorithm for generating genotypes from the super admixture model

To evaluate Algorithm 5, we assessed whether this algorithm is capable of generating genotypes that satisfy the moment constraints imposed by the super admixture model. More specifically, we examined if the estimated individual-level coancestry agrees with ***Q***′**Λ*Q*** and if the estimated coancestry among antecedent populations agrees with **Λ**.

To check these, we generated 300 unique combinations of (**Λ, *Q***) under the super admixture model as previously described. For each pair of (**Λ, *Q***), we (i) simulated ancestral allele frequencies ***a*** (*m* = 500, 000) by generating each *a*_*i*_ independently from Uniform(0, 1), (ii) generated genotypes ***X*** using Algorithm 5, (iii) estimated the individual-level coancestry 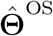 and (iv) applied Algorithm 1 to estimate 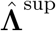 with 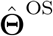 and ***Q*** as inputs. In (ii), a matrix of antecedent population allele frequencies ***P*** is generated at an intermediate step. We used both the double-admixture method and the NORTA method for generating ***P*** to compare their performances. For (iii), the OS estimate utilizes the minimum pairwise coancestry equal to 0, which might not hold here. We used the strategy described in Appendix B.1 to adapt the OS estimate to our simulation. We assessed the agreement between 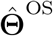 and **Θ** and between 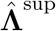 and **Λ** by 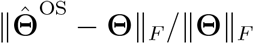 and 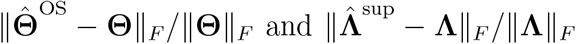, respectively. In Fig. C.3, we observed the majority of 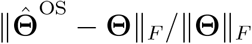 and 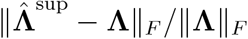 are less than 0.05 regardless of the method used for simulating ***P*** . These observations demonstrated our simulated genotypes satisfied the desired moment constraints.

**Figure C.3:**
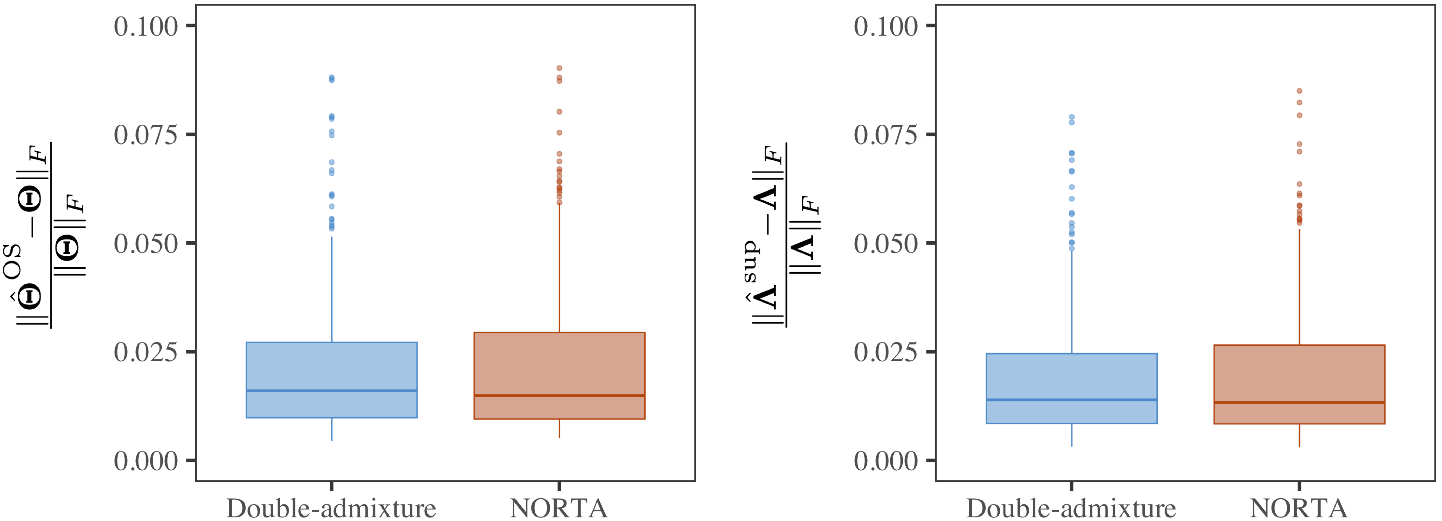
Box-plots of 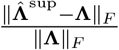 and 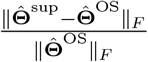 across 300 simulations.

#### C.6 Null *p*-value distribution of the hypothesis test of standard admixture versus super admixture

Recall the hypothesis test of the standard admixture model (null) versus the super admixture model (alternative):

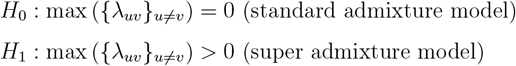

To check whether the true null hypothesis *p*-values calculated by Algorithm 7 are stochastically greater than or equal to the Uniform(0, 1) distribution, we generated 300 unique combinations of (**Λ, *Q***) under the standard admixture model. For each pair of (**Λ, *Q***), we (i) simulated ancestral allele frequencies ***a*** (*m* = 500, 000) by generating each *a*_*i*_ independently from Uniform(0, 1), (ii) generated genotypes ***X*** from the standard admixture model, (iii) applied Algorithm 7 to compute the *p*-values. We compared the empirical distribution of the *p*-values against Uniform(0, 1). Fig. C.4 shows that our proposed method is conservative, meaning it has a maximum type I error probability less than or equal to the nominal level of the test.

**Figure C.4:**
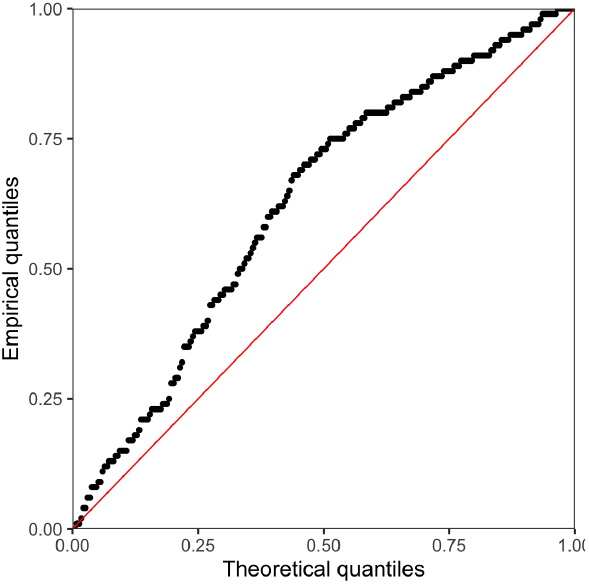
Quantile-quantile plots of true null *p*-values vs. Uniform(0, 1) distribution.

### D Supplementary analyses of human studies

#### D.1 Data processing

##### HO data

We downloaded the publicly available main Human Origins dataset and the Pacific dataset. The downloaded datasets are in the Eigensoft package format. We converted them to the PLINK format with the Eigensoft `convertf` function. We then merged the main Human Origins dataset with the Pacific dataset. These datasets have non-overlapping individuals that were genotyped using the same microarray platform. We excluded individuals from singleton subpopulations and the ancient individuals from the Lapita-Vanuatu population. We excluded SNPs with minor allele frequency (MAF) less than 0.01. The final dataset has 486, 981 SNPs and 2124 individuals.

##### HGDP data

We downloaded the publicly available HGDP dataset. We preserved loci that (i) are autosomal, biallelic SNPs, (ii) have MAF ≥ 0.01 and (iii) are in approximate linkage equilibrium with each other (*PLINK --indep-pairwise 1000kb 0*.*3*). The final dataset has 997, 431 SNPs and 929 individuals.

##### TGP data

We downloaded the publicly available TGP dataset. We preserved loci that (i) are autosomal, biallelic SNPs, (ii) are variants in the Yoruba individuals, (iii) have MAF ≥ 0.05 and (iv) are in approximate linkage equilibrium with each other (*PLINK --indep-pairwise 1000kb 0*.*3*). The final dataset has 712, 998 SNPs and 2583 individuals.

##### AMR subset of TGP

We identified individuals in the TGP dataset marked as `AMR` to create the AMR subset of TGP. We preserved loci that (i) are autosomal, biallelic SNPs, (ii) are variants in the Yoruba individuals, (iii) have MAF ≥ 0.01 and (iv) are in approximate linkage equilibrium with each other (*PLINK --indep-pairwise 1000kb 0*.*3*). The final dataset has 555, 145 SNPs and 353 individuals.

##### IND data

We obtained the Indian dataset from the authors of ref. [22]. We merged this dataset with the Central/South Asia population and the East Asia population of HGDP. These datasets have non-overlapping individuals that were genotyped using the same microarray platform. We excluded SNPs with MAF *<* 0.01 and SNPs with MAF differences greater than 0.2 to resolve the allele flipping issue. We also excluded SNPs with missing rates in IND or in the HGDP subset greater than 0.005. We applied this filter to keep high quality variants. The final dataset has 221, 499 SNPs and 698 individuals.

#### D.2 HGDP study analysis

We observed good concordance between the individual-level coancestry of HGDP for the OS and super admixture estimates (Fig. D.5) and early human migrations [29–32]. We note that the earliest major split occurred between Africa and MiddleEast from an out-of-Africa migration around 50 to 60 kya, resulting in the divergence between Sub-Saharan Africans and the remaining human populations. Another major split occurred between Central / South Asia and East Asia, revealing the separation between West Eurasians and East Asians around 40 to 45 kya. Among the East Asia clade, the Oceanians have the highest within subpopulation coancestry and lowest between subpopulation coancestry, consistent with the theory that Oceanians split earliest from the remaining East Asians.

**Figure D.5:**
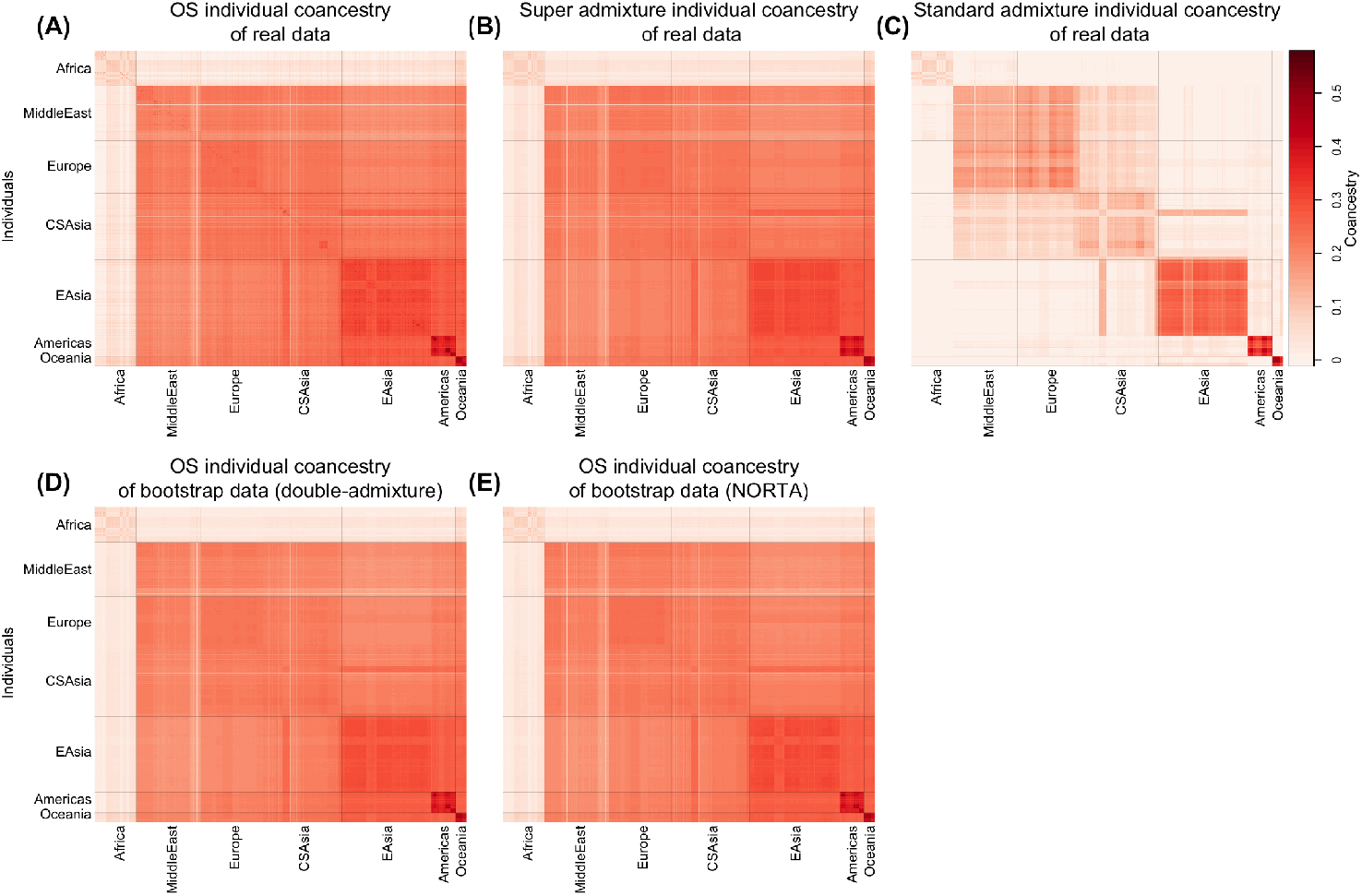
Heatmap of individual-level coancestry estimates in HGDP.

We confirmed that the super admixture antecedent population coancestry estimates are also compatible with known early human dispersals (Fig. D.6). Specifically, in Fig. D.6B the deepest split occurred roughly between individuals from Sub-Saharan Africa represented by the antecedent populations *S*_1_ and *S*_2_ and individuals outside of Sub-Saharan Africa represented by the remaining antecedent populations. Individuals outside of Sub-Saharan Africa further branched into two lineages: the West Eurasians represented by antecedent populations *S*_3_ and *S*_4_, and the East Asians represented by antecedent populations *S*_5_, *S*_6_, and *S*_7_. The Oceanians represented by *S*_7_ split from the majority of ancestral East Asians, while the remaining East Asians further diverged into present-day Asians (*S*_5_) and presentday Americans (*S*_6_).

**Figure D.6:**
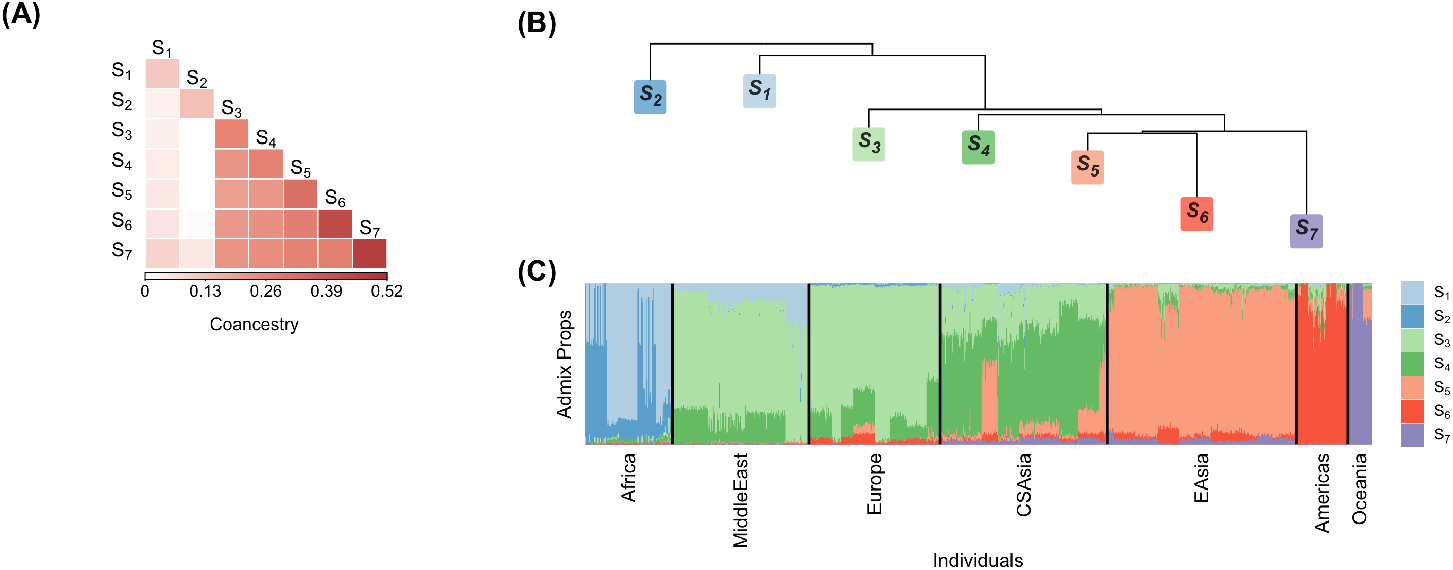
Heatmap of antecedent population coancestry estimates in HGDP. (B) Dendrogram representation of the antecedent population coancestry estimates. (C) Stacked bar plot of admixture proportions.

#### D.3 TGP study analysis

We also observed good correspondence between the individual-level coancestry OS and super admixture estimates of the TGP data (Fig. D.7) and known early human migrations [29–32]. Similarly to our analysis of HO and HGDP, the earliest major split is between AFR and the other populations, which reflects the divergence between Sub-Saharan Africans and the remaining of human populations. Another split occurs between EUR and EAS, revealing the separation between West Eurasians and East Asians.

As in our analysis of HO and HGDP, there is agreement between the estimated antecedent population coancestry and existing results. In Fig. D.8B, the deepest split occurred roughly between individuals from Sub-Saharan Africa represented by antecedent population *S*_1_ and individuals outside of Sub-Saharan Africa represented by the rest of the antecedent populations. We also noted the divergence between the Europeans represented by antecedent population *S*_3_, and the Asians represented by antecedent populations *S*_2_, *S*_4_ and *S*_5_. The Americans sampled in TGP appear to have a higher European ancestry compared to that in the HO and HGDP datasets.

**Figure D.7:**
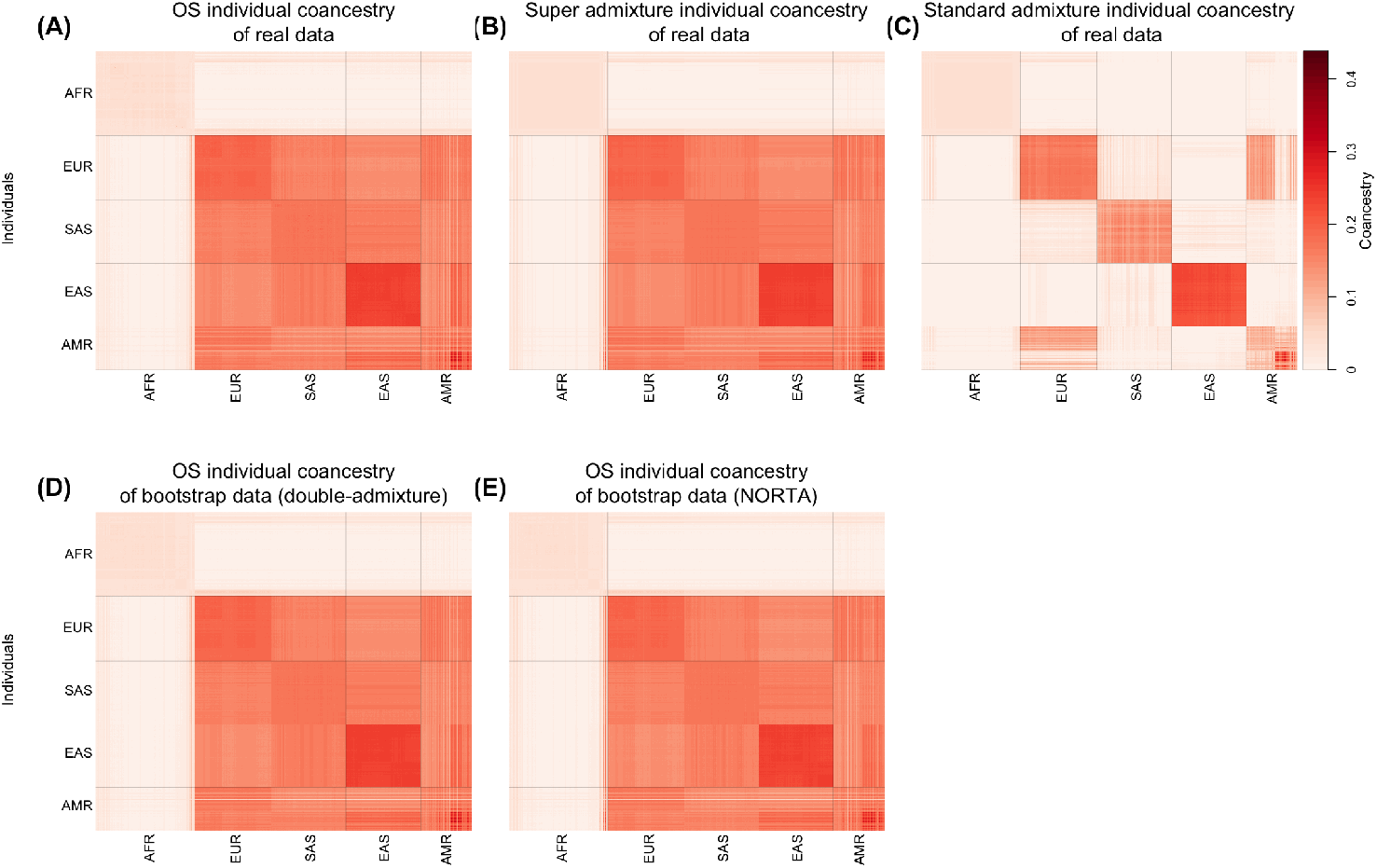
Heatmap of individual-level coancestry estimates in TGP.

**Figure D.8:**
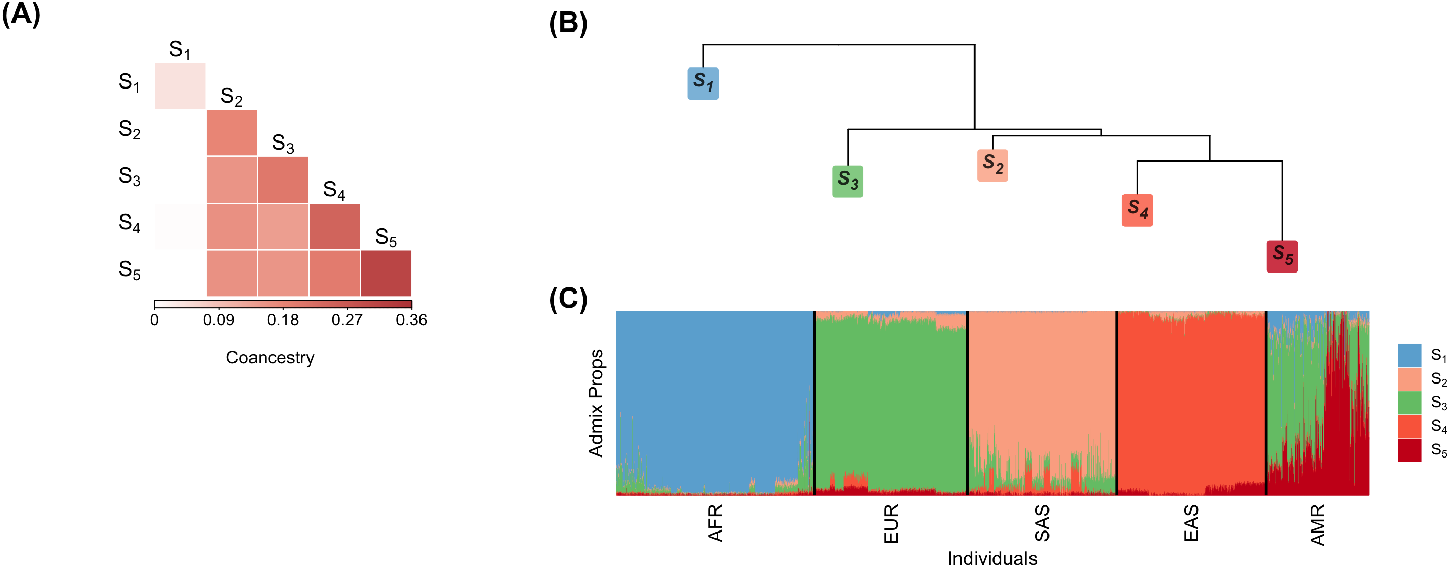
Heatmap of antecedent population coancestry estimates in TGP. (B) Dendrogram representation of the antecedent population coancestry estimates. (C) Stacked bar plot of admixture proportions.

#### D.4 Confirming significant hypothesis tests of standard admixture versus super admixture in the human studies

We applied Algorithm 7 to each of the five human datasets to statistically evaluate the presence of coancestry among antecedent populations. Each panel of Fig. D.9 shows the distribution of the *B* = 1000 bootstrap null test-statistics for each dataset. The observed test-statistic *U*_obs_ for each data set is noted on the top-right of each panel. In each data set *U*_obs_ exceeded all bootstrap null test-statistics, implying *p*-value *<* 0.001 for each.

#### D.5 Comparing the individual-level coancestry estimates

In Table D.1, we compared 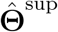 and 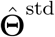 to the general OS estimate of individual-level coancestry 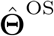 from ref. [4] on the five data sets. The super admixture coancestry estimate has about 10 to 40 times smaller distance to the OS estimate compared to the standard admixture estimate.

#### D.6 Selecting the number of antecedent populations

We utilized the structural Hardy-Weinberg (sHWE) framework [12] for determining the number of antecedent populations *K*, as outlined in that work. The approach considers a range of *K* values for a model of structure that results in estimated IAFs, which is the case for our framework. For each *K*, a hypothesis test is performed for each SNP of the assumption that *x*_*ij*_|*π*_*ij*_ ∼ Binomial(2, *π*_*ij*_) (*j* = 1, 2, … , *n*) based on the estimates 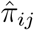 and a goodness-of-fit statistic with a parametric bootstrap null distribution. This results in *m* p-values per value of *K*.

As proposed in the sHWE framework, for each value of *K*, we (i) calculated the *m* sHWE *p*-values, (ii) binned the sHWE *p*-values into equal-sized bins (number of bins, *C* = 150), (iii) removed the first bin [0, 1*/C*), and (iv) calculated the following negative entropy that measures how well the sHWE *p*-values follow the Uniform(0, 1) distribution,

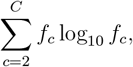

where *f*_*c*_ denotes the proportion of *p*-values in the *c*-th bin.

**Figure D.9:**
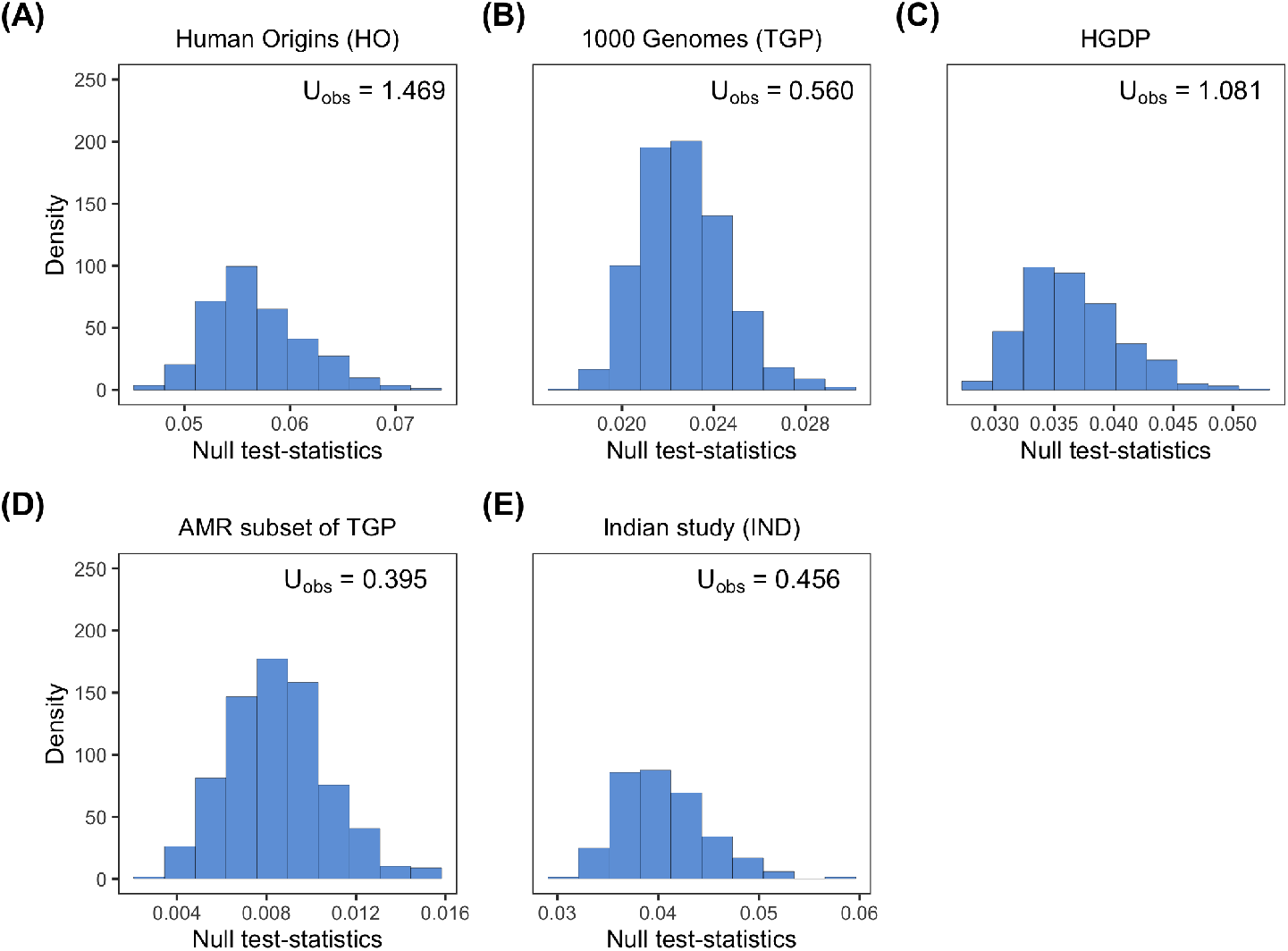
Distributions of test-statistics for the hypothesis tests of standard admixture versus super admixture across all five human studies.

**Table D.1:** The distance between 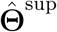 and 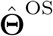 and between 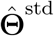 and 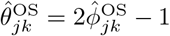 for each of the five data sets.

| | $\frac{1}{n} \ \hat{\Theta}^{\text{sup}} - \hat{\Theta}^{\text{OS}}\ _F$ | $\frac{1}{n} \ \hat{\Theta}^{\text{std}} - \hat{\Theta}^{\text{OS}}\ _F$ |
| --- | --- | --- |
| HO | 0.003 | 0.114 |
| AMR | 0.006 | 0.124 |
| IND | 0.006 | 0.052 |
| HGDP | 0.008 | 0.146 |
| TGP | 0.002 | 0.087 |
| | $\ \hat{\Theta}^{\text{sup}} - \hat{\Theta}^{\text{OS}}\ _F / \ \hat{\Theta}^{\text{OS}}\ _F$ | $\ \hat{\Theta}^{\text{std}} - \hat{\Theta}^{\text{OS}}\ _F / \ \hat{\Theta}^{\text{OS}}\ _F$ |
| HO | 0.021 | 0.751 |
| AMR | 0.023 | 0.507 |
| IND | 0.073 | 0.613 |
| HGDP | 0.040 | 0.724 |
| TGP | 0.017 | 0.700 |

We determined *K* to be the value achieving the minimum negative entropy. In the event of a plateau where the entropy is more or less the same over a range of *K*, we chose a small value of *K*, where the plateau began. This resulted in *K* = 11 for HO, *K* = 7 for HGDP, *K* = 5 for TGP, *K* = 7 for IND, and *K* = 3 for AMR.

In Fig. D.10, we observed a consistent decline in 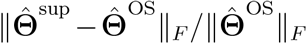 as *K* increased. In Fig. D.11, we noted an increase in 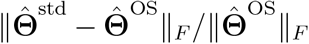 as *K* increased. This implies the standard admixture fit cannot be improved with a larger *K*. In the following two subsections, we show that the results for HO and IND are similar over a range of *K* close to the values that we selected.

**Figure D.10:**
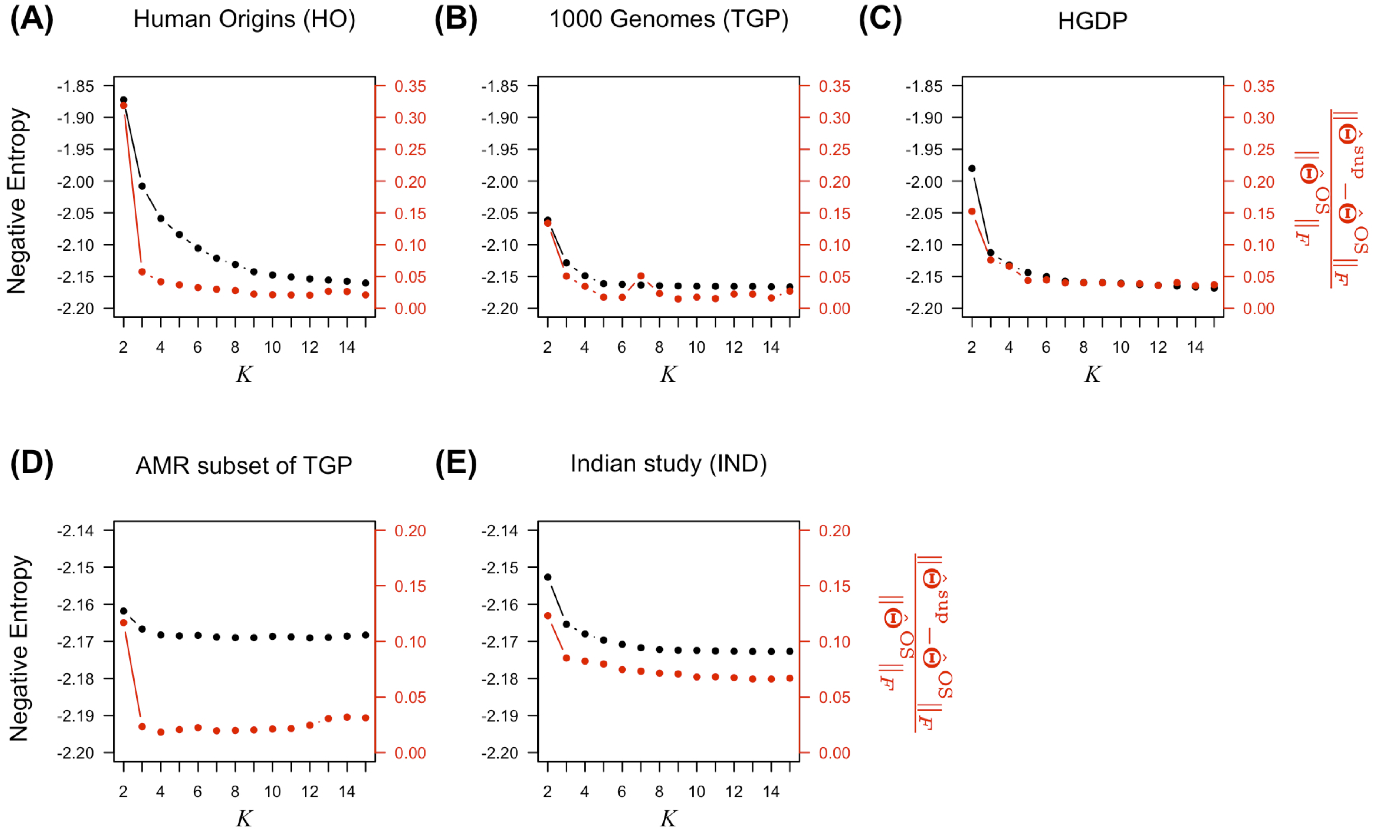
The negative entropy and 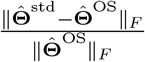 across different numbers of antecedent populations.

**Figure D.11:**
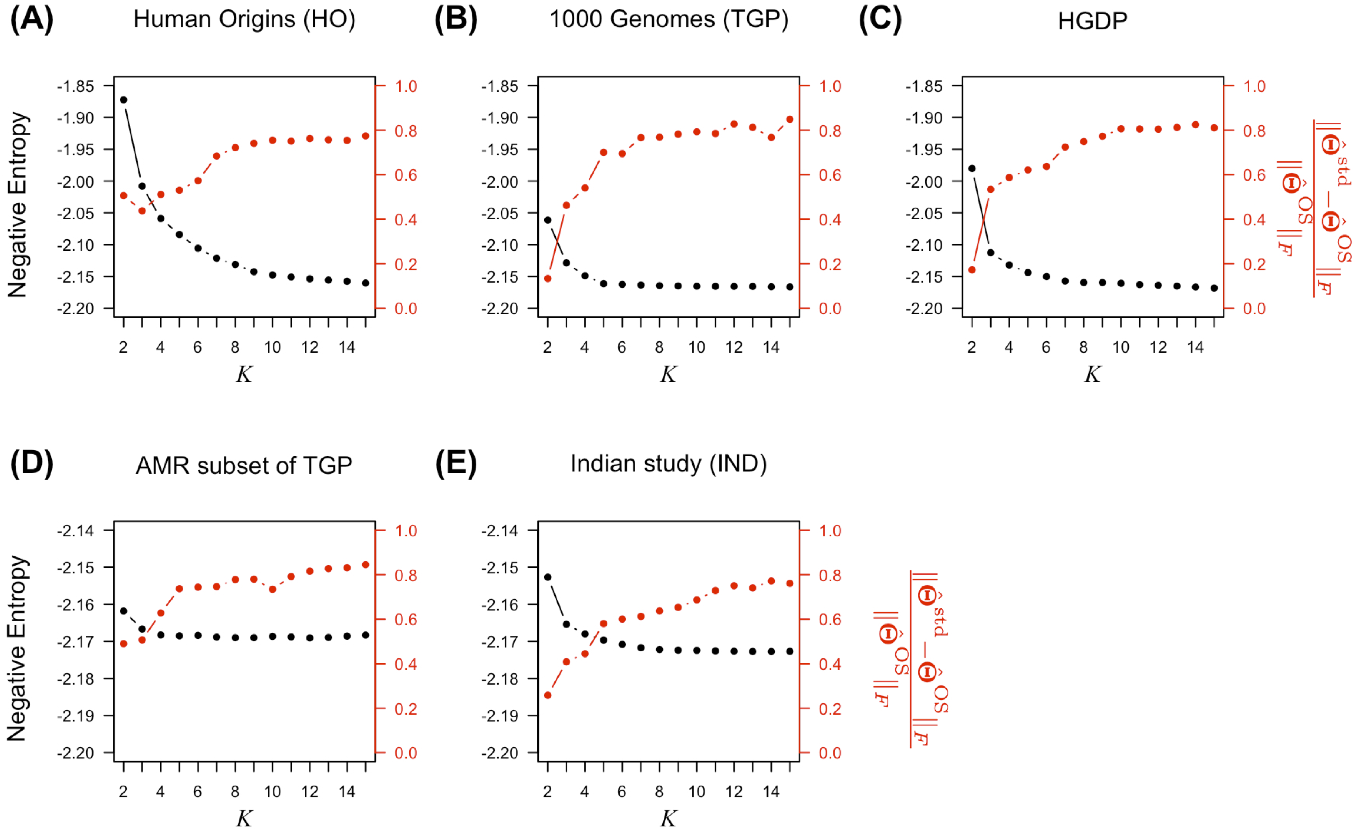
The negative entropy and 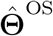 across different numbers of antecedent populations.

#### D.7 Analysis of HO over a range of antecedent population numbers

In our analysis of HO, we utilized *K* = 11 antecedent populations. We also analyzed the HO dataset for *K* = 7, 8, 9, 10 in this section to demonstrate the choice of *K* did not have a major impact on our results and conclusions. In Figs. D.12 to D.15, we observed the estimated antecedent population coancestries and the admixture proportions were consistent for *K* = 7, 8, 9, 10.

**Figure D.12:**
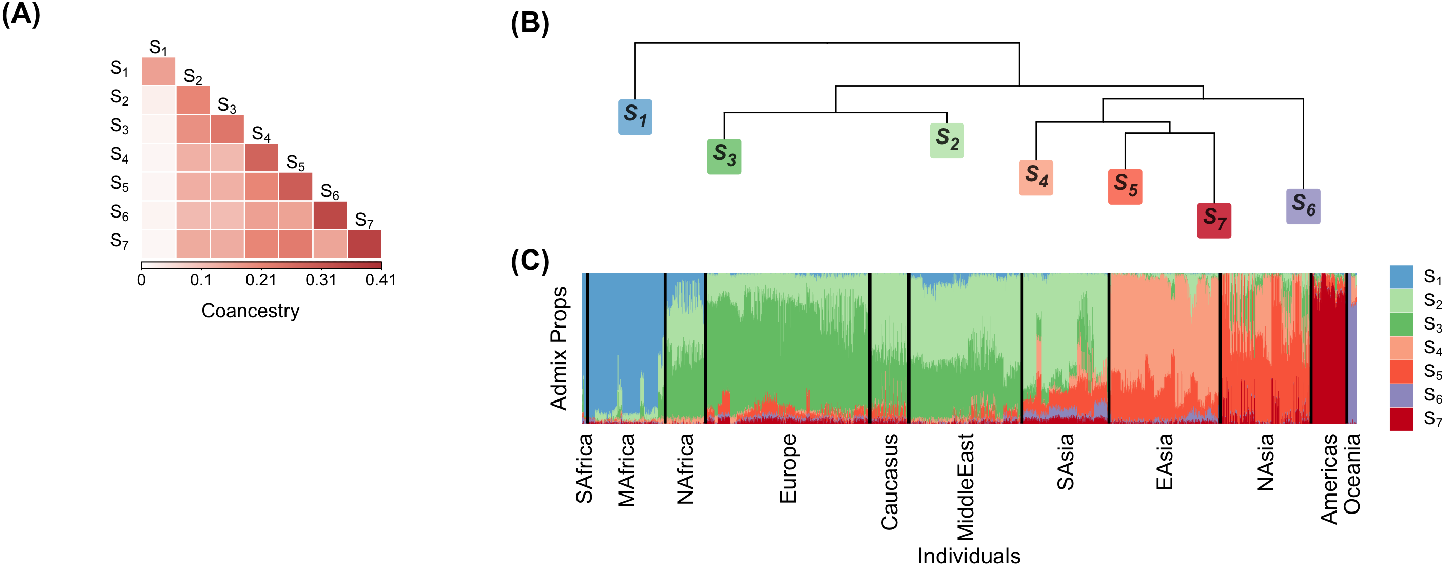
(A) Heatmap of antecedent population coancestry estimates in HO with *K* = 7. (B) Dendrogram representation of the antecedent population coancestry estimates. (C) Stacked bar plot of admixture proportions.

**Figure D.13:**
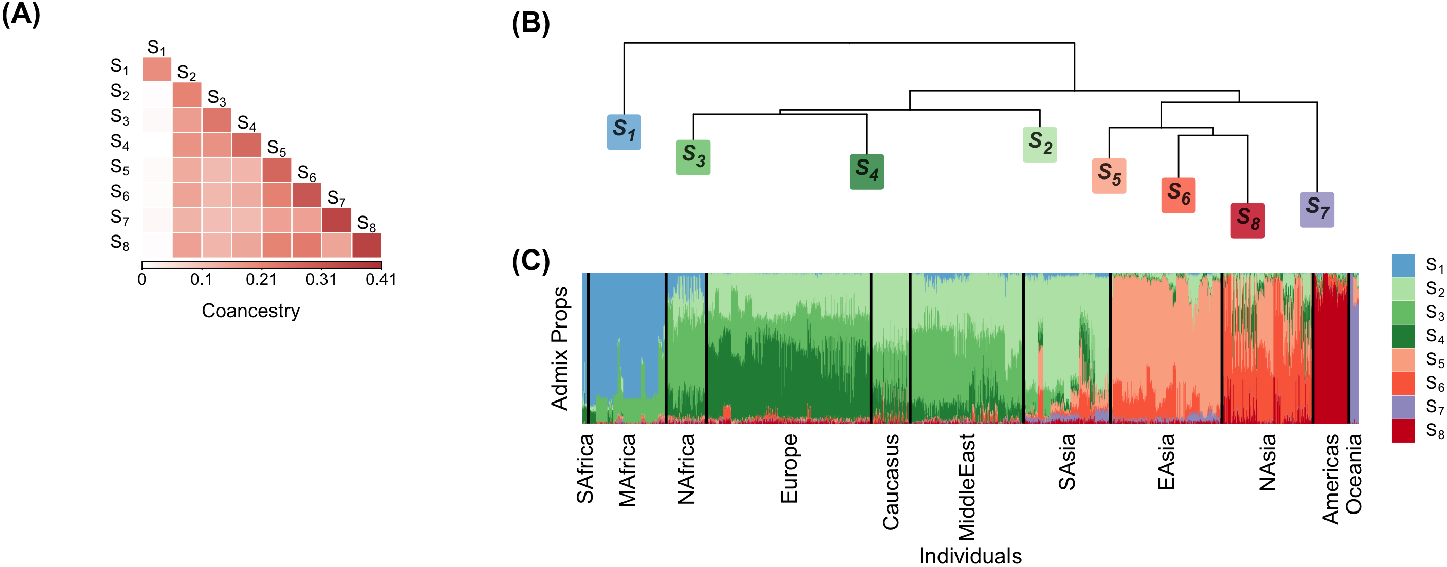
(A) Heatmap of antecedent population coancestry estimates in HO with *K* = 8. (B) Dendrogram representation of the antecedent population coancestry estimates. (C) Stacked bar plot of admixture proportions.

**Figure D.14:**
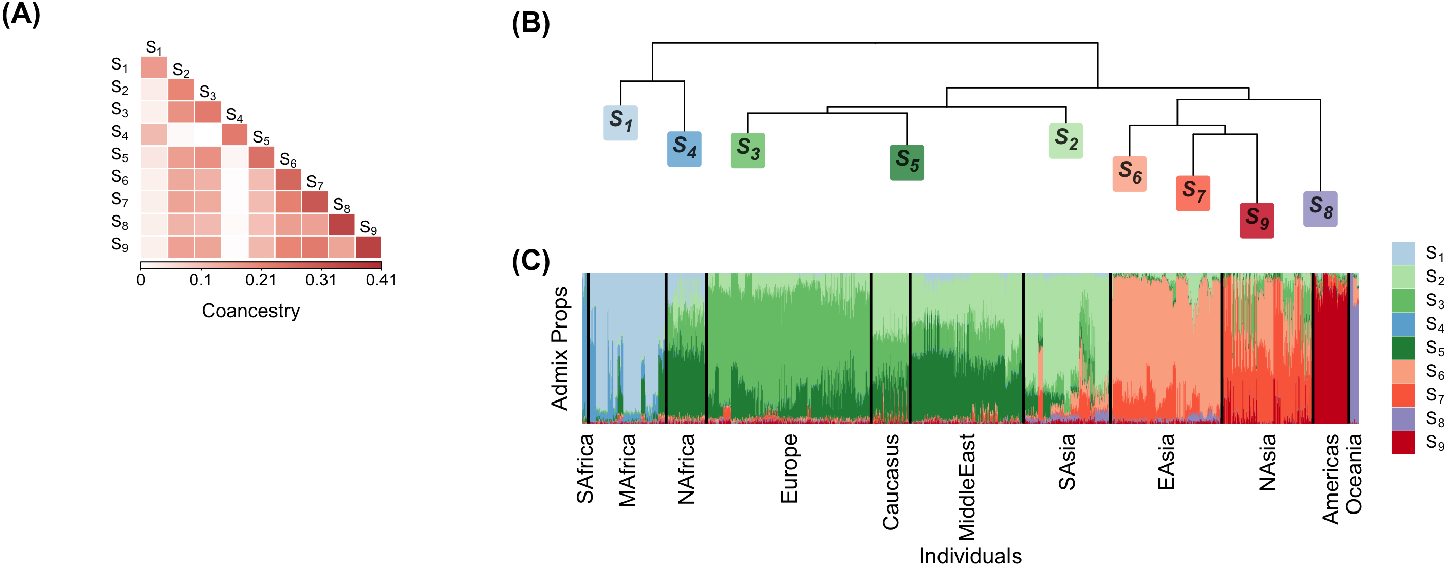
(A) Heatmap of antecedent population coancestry estimates in HO with *K* = 9. (B) Dendrogram representation of the antecedent population coancestry estimates. (C) Stacked bar plot of admixture proportions.

**Figure D.15:**
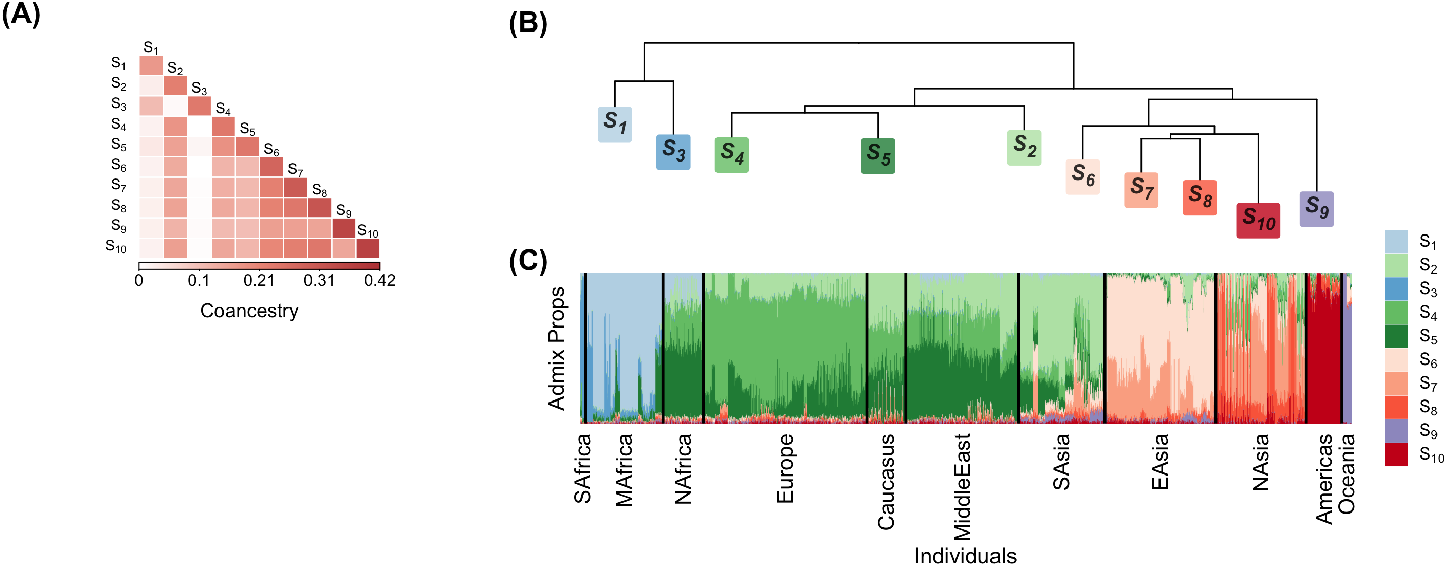
(A) Heatmap of antecedent population coancestry estimates in HO with *K* = 10. (B) Dendrogram representation of the antecedent population coancestry estimates. (C) Stacked bar plot of admixture proportions.

#### D.8 Analysis of IND over a range of antecedent population numbers

In our analysis of IND, we utilized *K* = 7 antecedent populations. In Figs. D.16 to D.19, we observe the estimated antecedent population coancestries and the admixture proportions were consistent for *K* = 3, 4, 5, 6.

**Figure D.16:**
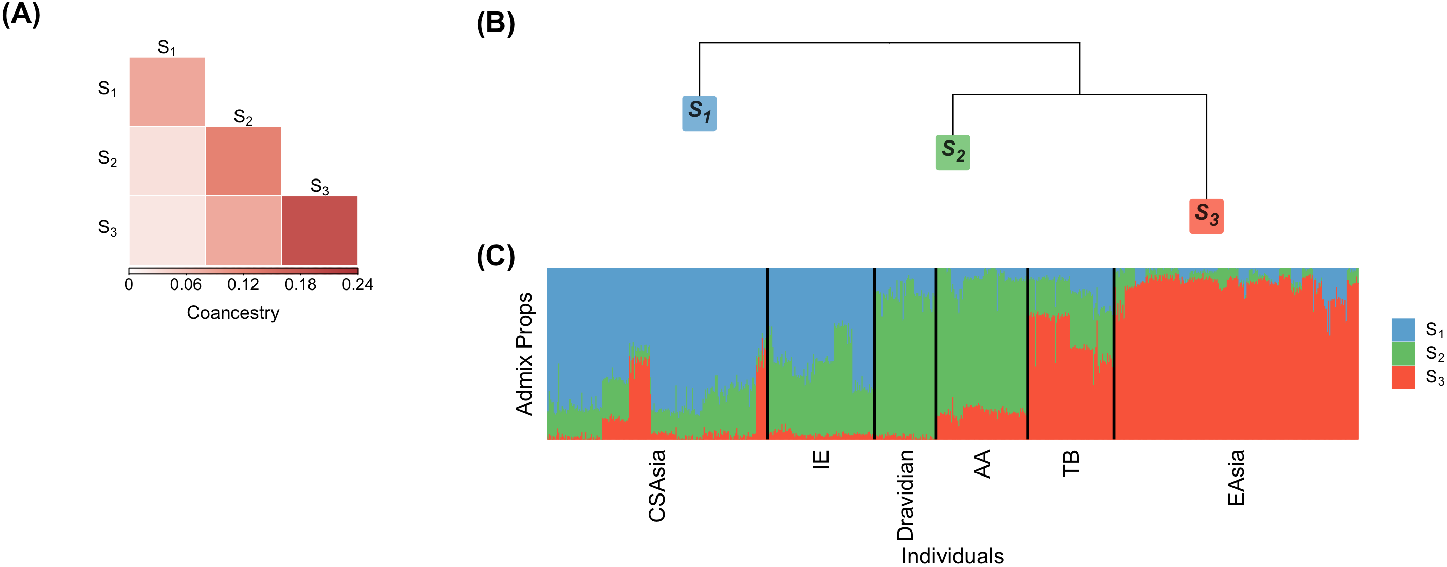
(A) Heatmap of antecedent population coancestry estimates in the merged data sets of IND with Central/South Asians and East Asians of HGDP with *K* = 3. (B) Dendrogram representation of the antecedent population coancestry estimates. (C) Stacked bar plot of admixture proportions.

**Figure D.17:**
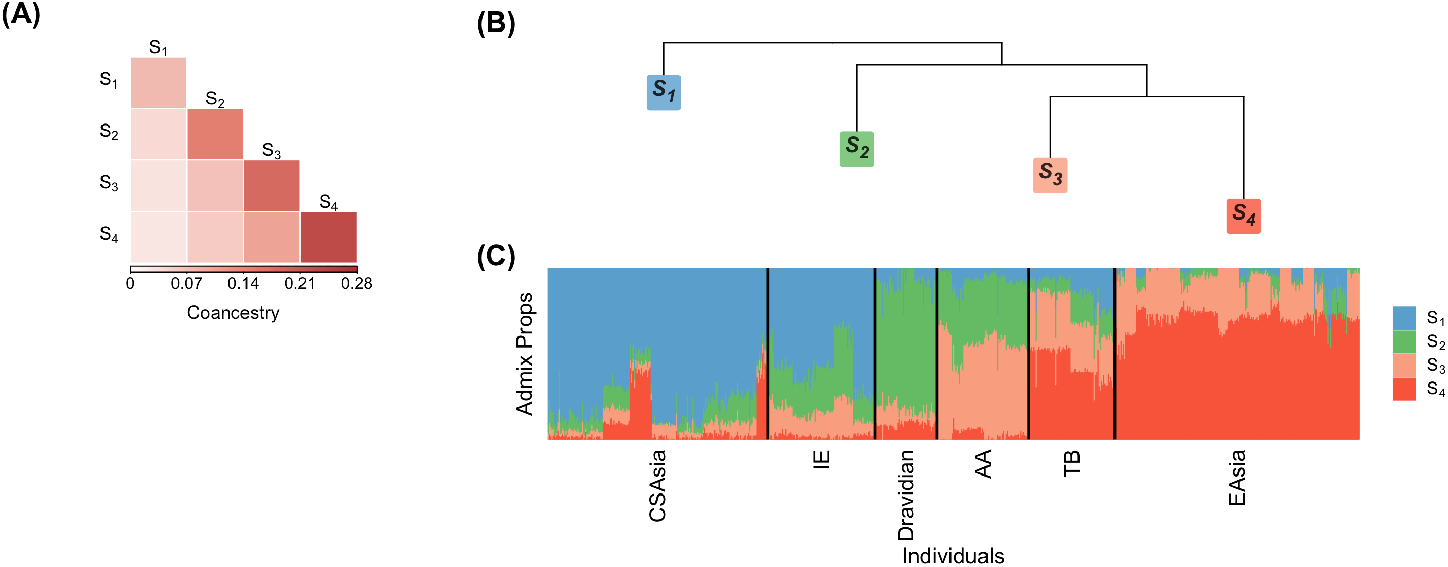
(A) Heatmap of antecedent population coancestry estimates in the merged data sets of IND with Central/South Asians and East Asians of HGDP with *K* = 4. (B) Dendrogram representation of the antecedent population coancestry estimates. (C) Stacked bar plot of admixture proportions.

**Figure D.18:**
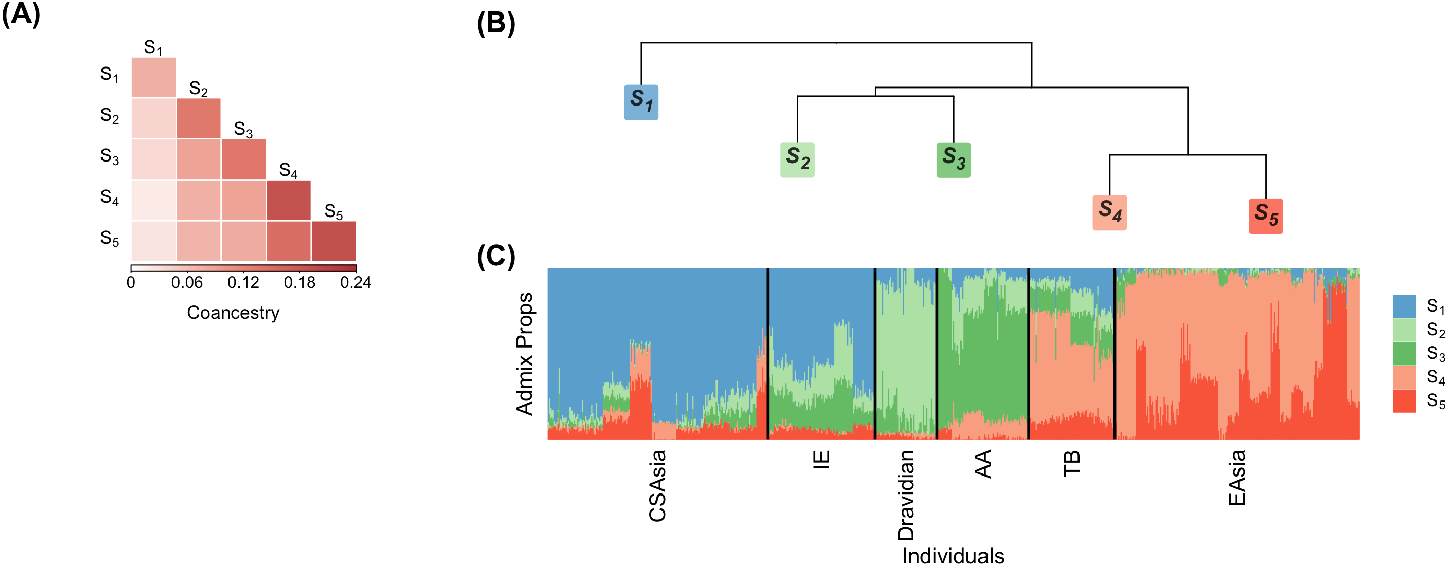
(A) Heatmap of antecedent population coancestry estimates in the merged data sets of IND with Central/South Asians and East Asians of HGDP with *K* = 5. (B) Dendrogram representation of the antecedent population coancestry estimates. (C) Stacked bar plot of admixture proportions.

**Figure D.19:**
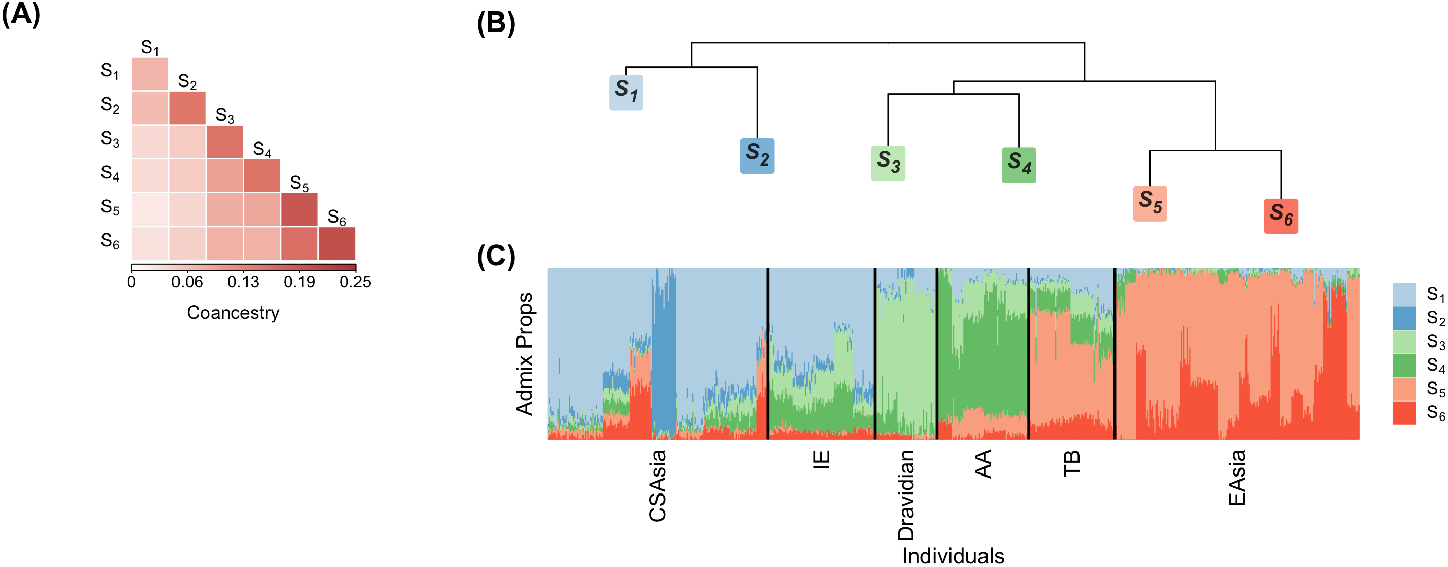
(A) Heatmap of antecedent population coancestry estimates in the merged data sets of IND with Central/South Asians and East Asians of HGDP with *K* = 6. (B) Dendrogram representation of the antecedent population coancestry estimates. (C) Stacked bar plot of admixture proportions.

## Footnotes

1 Note also that the OS estimate of **Θ** is equal to the OS estimate of kinship, **Φ**, except for the diagonal elements where 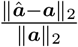, as shown in Eq. (3).

